# *FindingNemo*: A Toolkit for DNA Extraction, Library Preparation and Purification for Ultra Long Nanopore Sequencing

**DOI:** 10.1101/2024.08.16.608306

**Authors:** Inswasti Cahyani, John Tyson, Nadine Holmes, Josh Quick, Chris Moore, Nick Loman, Matthew Loose

## Abstract

Since the advent of long read sequencing, achieving longer read lengths has been a key goal for many users. Ultra-long read sets (N50 ≥ 100 kb) produced from Oxford Nanopore sequencers have improved genome assemblies in recent years. However, despite progress in extraction protocols and library preparation methods, ultra-long sequencing remains challenging for many sample types. Here we compare various methods and introduce the *FindingNemo* protocol that: (1) optimises ultra-high molecular weight (UHMW) DNA extraction and library clean-up by using glass beads and Hexamminecobalt(III) chloride (CoHex), (2) can deliver high ultra-long sequencing yield of >20 Gb of reads from a single MinION flow cell or >100 Gb from PromethION devices (R9.4 to R10.4 pore variants), and (3) is scalable to using fewer input cells or lower DNA amounts, with extraction to sequencing possible in a single working day. By comparison, we show this protocol is superior to previous ones due to precise determination of input DNA quantity and quality by cell count, sample dilution and homogenization approaches.

## Introduction

The first ultra-long (UL) sequencing protocols on nanopore sequencing instruments resulted in highly contiguous human genome assemblies (Jain et al. 2018a; Quick 2018). Later methods improved on these assemblies through increased yields (Logsdon et al. 2021; Miga et al. 2020). Oxford Nanopore Technologies (ONT) subsequently launched UL sequencing kits, starting with ULK001 on R9.4 flow cells and more recently ULK114 on R10.4.1 flow cells. The ULK001 protocol utilised extraction and clean-up steps using silicon coated discs from Circulomics Inc. (Pacific Biosciences, California, USA), which was subsequently replaced with the ONT star-shaped matrix for library precipitation in the ULK114 kit. These UL kits introduced a range of protocol improvements both in library preparation and sequencing performance to maximise read length and generated reads with an N50 > 100kb from both the MinION and PromethION flow cells.

In our experience, we have found that the most critical step in obtaining high throughput, high occupancy UL sequencing libraries is the DNA fragmentation step, followed by the final clean-up step prior to loading the library. The UL protocol uses a transposase enzyme complex (transposome) to fragment DNA. During fragmentation, the DNA should be homogeneous and accessible to this fast-acting transposome. Two methods are effective in reducing the number of transposase cuts to increase read length: either using a high concentration of high molecular weight (HMW) DNA as shown by Jain et al in sequencing the human genome (Jain et al. 2018a) or diluting the transposase relative to DNA (Jain et al. 2018b). UL libraries are difficult to clean up using conventional magnetic solid phase reversible immobilisation (SPRI) approaches as the DNA can be hard to elute and so sheared during this; SPRI beads are mostly used for PCR clean-up where the fragment length is much shorter (Rodrigue et.al 2010; Stortchevoi et.al. 2020). In addition, the library is a complex of motor protein and DNA, and so alcohol-based precipitation and washes are not compatible. In our search for suitable alternative approaches, we found that hexamine cobalt (III) (CoHex) cations stabilise and condense DNA and reasoned they may enable alcohol-free precipitation of nanopore sequencing libraries (Allers and Lichten 2000; Kankia et al. 2001).

Early versions of the ONT ultra-long (UL) protocol required extraction and library clean-up starting from 6 million human cells, or an equivalent input amount of DNA (PacBio 2023, Methods). The complete extraction and library preparation protocol takes between two to three days in the laboratory. However, different sample types often require tailored extractions to provide optimal DNA for sequencing, so we sought to use a wide range of sample extraction protocols to obtain UL reads on ONT platforms. We also explored reducing both sample input requirements and the total time for library preparation for sequencing scalability.

Here we describe the *FindingNemo* protocol (named after the characteristic orange colour of the CoHex buffer) for the generation of high occupancy ultra-long reads on nanopore platforms. This protocol can generate equivalent or more throughput to disc-based methods and may have additional advantages in tissues and non-human cell material. The protocol can also be tuned to enable extraction from as few as one million human cell equivalents or 5 μg of human ultra-high molecular weight (UHMW) DNA as input and enables extraction to sequencing in one working day. The clean-up method can be used to generate ultra-long libraries from DNA extracted with phenol chloroform, SDS lysis and CTAB approaches as well as commercial kits such as the NEB Monarch HMW gDNA kits and those from Circulomics (now Pacific Biosciences). Oxford Nanopore Technologies released similar protocols using a spermine-based precipitation buffer (PPT) instead of CoHex and no DNA binding substrate (ONT, ULK001) and then incorporating a star (ONT, ULK114) after our initial protocol release on protocols.io (Cahyani 2021a). We compare all these methods here.

## Results

### A. The *FindingNemo* protocol compiles and optimises available ultra-long sequencing protocols

#### Previously developed protocols

Both the standard and ultra-long (UL) sequencing workflows consist of three main steps: DNA extraction, library preparation, and library clean-up prior to flow cell loading (Figure 1a). Ultra-long sequencing necessitates minimal DNA shearing throughout the library preparation process and during the extraction methods to maintain ultra-high molecular weight (UHMW) DNA. Early experiments pioneering UL sequencing used the transposase-based rapid kit (ONT, e.g., RAD004) to obtain reads with an N50 greater than 100 kb (Jain et al. 2018a). This method eliminated the need for DNA shearing before library preparation. However, the yield from rapid runs was typically low, which might be a consequence of the protocol itself. The lack of purification or clean-up of rapid libraries prior to flow cell loading likely contributed to reduced yields compared to ligation-based sequencing approaches. To better understand the complex interplay of DNA quality, library preparation, read length, and yield, we tested a range of UL protocols aimed at maximising both read length and yield (Figure 1b).

**Figure 1.**
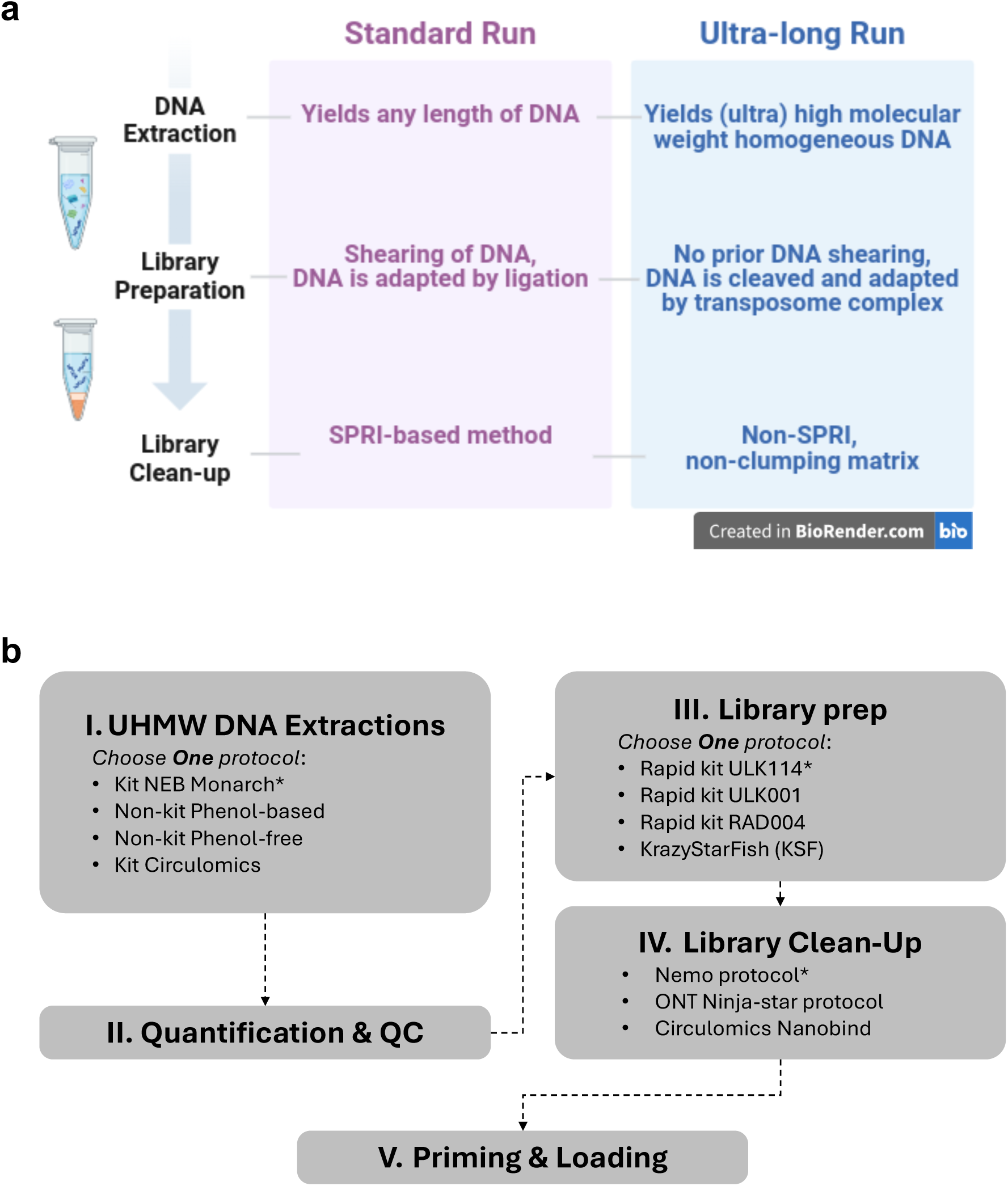
Summary of the FindingNemo toolkit: (a) Similarities and differences between the standard and ultra-long (UL) protocol that centres on the way UHMW DNA is handled. (b) The route of FindingNemo toolkit consists of five main steps, with options at the extraction, library preparation and clean-up steps. Options with asterisks (*) are our current lab workflow for the ultra-long protocol.

The goal in obtaining UL reads with the rapid kit is to ensure that each DNA molecule is cut only once by a transposase complex, maximising read length. Initially, Quick et al achieved this by saturating the transposase reaction with UHMW DNA (Quick 2018) (Table 1A). An alternative approach by Logsdon et al. was to reduce the amount of transposase for a fixed amount of DNA (Logsdon 2021) (Table 1B). The latter method typically results in sub-optimal pore occupancy (i.e. the number of available pores sequencing at any time) due to the lower number of adapted DNA ends available for sequencing because of the reduced amount of transposase. In testing the Quick protocol, we observed two different read length distributions (see Supplemental Figure S1a, b). Whilst both had N50s of approximately 90 kb, one was dominated by shorter reads. This suggests that the DNA was not cut uniformly. Moreover, although these protocols generated UL reads, they suffered from high variability in output N50, occupancy and yield (Supplemental Figure S1a, b). Typically, as N50 increases, yield and occupancy drop (Supplemental Table S1A, B).

**Table 1.** Compilation of ultra-long (UL) library preparation protocols, optimized in human GM12878 cells.

|  | Previous Protocols |  |  |  | New Protocols |  |  |  |  |
| --- | --- | --- | --- | --- | --- | --- | --- | --- | --- |
|  | Quick's UL<br>(Quick, 2018) | Logsdon's<br>UL<br>(Logsdon,<br>2020) | Rocky<br>Mountain<br>(Tyson, 2020) | Krazy StarFish<br>(KSF) | ULK001 | RAD004-Nemo | ULK001-Nemo | ULK114 | ULK114-Nemo |
|  | A | B | C | D | E | F | G | H | I |
| Input cell number* | 50 million | 20-70 million | >2 million | >2 million | 6 million | 1-6 million | 1-6 million | 6 million | 6 million |
| Input DNA amount | 15 µg | 2-3 µg | ≥7.5 µg | ≥7.5 µg | 40 µg | 5-40 µg | 5-40 µg | 40 µg | 40 µg |
| ONT Library preparation kit | RAD004 | RAD004 | LSK109 | RAD004 | ULK001 | RAD004 | ULK001 | ULK114 | ULK114 |
| Fragmentation Volume | Small<br>(15 ml) | Small<br>(18 ml) | Medium<br>(60-100 ml) | Medium<br>(2x 100 ml) | Large<br>(1ml) | Large<br>(0.5-1 ml) | Large<br>(0.5-1 ml) | Large<br>(1ml) | Large<br>(1-1.2 ml) |
| Library clean-up protocol | No | No | No | Partial using<br>filter paper | Nanobind<br>(Circulomics) | Nemo | Nemo | No | Nemo |
| Processing time (excluding incubations) | 15 min | 2.5 hours | 18-24 hours | 30 min | ~2.5 hours | ~2.5 hours | ~2.5 hours | ~2.5 hours | ~2.5 hours |
| Input library per MinION flow cell | 10-15 µg | 2-3 µg | 1 to 3 µg | 2 to 4 µg | 5 to 6 µg | 1 to 5 µg | 1 to 5 µg | NA | NA |
| No. of loads per library (MinION) | 1 | 1 | 1 to 3 | 3 | 6 | 1 to 6 | 1 to 6 | NA | NA |
| Expected yield per MinION flow cell | ~3 Gb | 1-2 Gb | 10 to >20 Gb<br>(depends on<br>N50) | 6-8 Gb<br>(3 loads) | >20 Gb<br>(3 loads) | >15 Gb<br>(3 loads) | ~20 Gb<br>(3 loads) | NA | NA |
| Input library per PromethION flow cell | NA | NA | NA | NA | 10-13 µg | 10-13 µg | 10-13 µg | 10-13 µg | 10-13 µg |
| No. of loads per library (PromethION) | NA | NA | NA | NA | 3 | 1 to 3 | 1 to 3 | 3 | 3 to 4 |
| Expected yield per PromethION flow cell (Gb) | NA | NA | NA | NA | NA | NA | 24 Gb<br>(1 load) | ~85 Gb | up to<br>>100 Gb |
| Ultra-long read length (≥100 kb) | Yes | Yes | No | Yes | Yes | Yes | Yes | Yes | Yes |

(\*) cell number was dependent on the extraction protocol used; here it is listed as the number of cells required by the original protocol; if cell number is unknown, use the DNA amount; (NA) not available

The Rocky Mountain protocol (Tyson, 2020) (Table 1C) was developed to use the ligation kit (ONT, e.g. LSK109) to address the yield issues experienced with the rapid kit. This protocol utilised varying concentrations of salts and polyethylene glycol (PEG) to precipitate longer reads after the library preparation step. The ligation kit requires light shearing of the input DNA, resulting in non-UL N50s albeit with much improved yields (data not shown). Therefore, this protocol was further modified to use the rapid kit followed by precipitation of the tagmented DNA using a filter paper disc or star as a form of SPRI matrix (KrazyStarFish; Table 1D). The KrazyStarFish (KSF) protocol worked consistently in producing UL N50s, but occupancies and yields were still suboptimal, perhaps a consequence of having DNA clean-up prior to sequencing adapter ligation, consequently producing many free adapters (Supplemental Figure S1c, d; Supplemental Table S1C, D; Methods).

#### New and developing protocols

ONT released a rapid kit for UL sequencing, SQK-ULK001 (hereafter ULK001) in which the transposase reaction is performed in a large volume (Table 1E), keeping the DNA concentration at approximately 50 ng/μl. We hypothesised that larger reaction volumes facilitated even diffusion and mixing of DNA and transposase complexes in a more homogeneous state in solution, a feature that we included in our Nemo protocols (Table 1F, G). Efficient transposase reactions result in consistent UL sequencing output as shown by the high rates of occupancy (Supplemental Figure S1e-g, S2, Figure 2, and Supplemental Table S1). Occupancy as a parameter is specifically defined by the percentage of sequencing pores (i.e. pores in ‘adapter’ and ‘strand’ states) compared to all pores in available states (i.e. ‘adapter’, ‘strand’ and ‘pore’ state).

**Figure 2.**
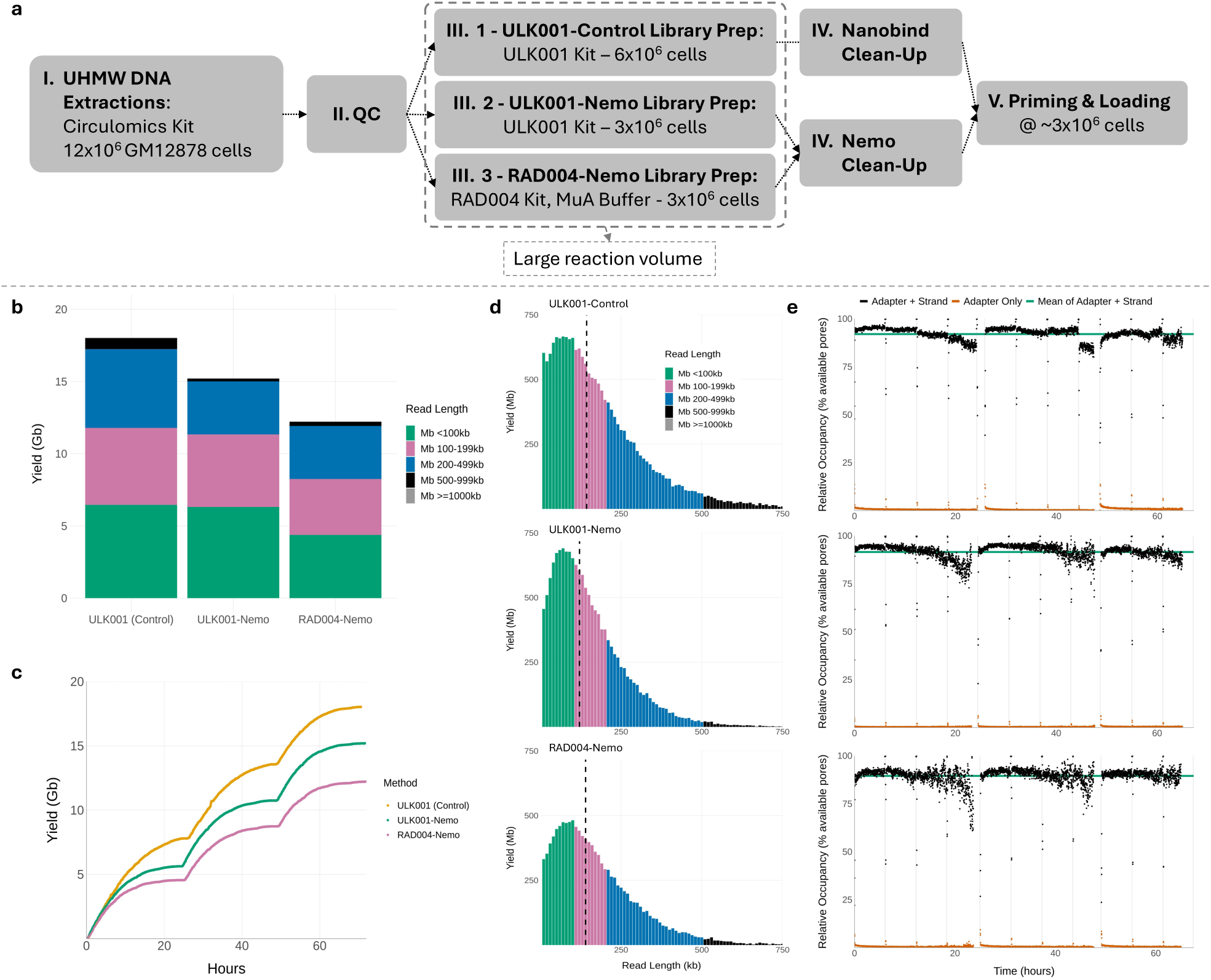
Validation of the FindingNemo protocol - sequencing outputs between rapid-based library preparation protocols (ULK001-Nemo and RAD004-Nemo protocol) showing comparable performance when compared to the control ULK001 protocol. (a) Workflow of the library preparation protocols, (b) Total yields sub categorised by read lengths, (c) Time lapse of yields, (d) Read length distributions; dashed black vertical lines denote N50s, (e) Time lapse of relative pore occupancies. Each library was loaded three times (i.e. total DNA equivalent to 3 million GM12878 cells) on a MinION R.9.4.1 flow cell following a nuclease flush protocol. All data are after 65 hours of sequencing on the GridION platform.

We next tested the performance of RAD004 and ULK001 rapid-based kit protocols, modifying parameters such as the number of input cells, the RAD004 dilution buffer, and the clean-up steps whilst maintaining the large volume ratio, and compared these to the control ULK001 protocol (Figure 2a). Regardless of clean-up method (Nanobind disc or glass beads), ULK001-based protocol outputs were comparable in terms of N50s and occupancies (Figure 2b-e). Meanwhile, the output of the library prepared with the RAD004 protocol showed slightly lower performance in general (Figure 2b-e and Supplemental Table S1G).

The total yield of an UL sequencing is not a direct measure of the library quality as yield is also flow cell dependent. The control ULK001 library produced the highest total yield (Figure 2b-c) likely because of the highest pore count of the flow cell (Supplemental Table S1E, Supplemental Figure S3a). Normalising yields between flow cells by taking a ratio of yield to the number of pores utilised during a period of data collection (i.e. yield per pore) allows comparison of run performance. When normalised, the yields per pore of the two ULK001 libraries were comparable (Supplemental Table S1E, F). In all three libraries, the distribution of read lengths against read quality were similar (Supplemental Figure S3b), as well as sequencing speed (Supplemental Figure S3c). These similarities imply that either of the rapid-based kits could be used to successfully produce UL sequencing outputs. Considering all sequencing output parameters, the ULK001 was the optimal protocol for UL sequencing compared to the RAD004 (Supplemental Table S2).

### B. The *FindingNemo* protocol efficiently purifies and accurately quantifies UHMW DNA and UL libraries

#### DNA precipitation and recovery: maintaining quality and homogeneity

Prior to the launch of the ULK001 kit we developed a protocol to clean the library after adapter ligation using DNA precipitation chemistry compatible with the motor protein-DNA complex. The filter paper from the KSF protocol was replaced with glass beads, adapting the glass bead’s effectiveness to act as a DNA-binding substrate as in the Monarch extraction kit protocol (NEB, T3050). We did not observe a significant difference in DNA recovery when using borosilicate glass beads compared to other common types of glass (data not shown). We surveyed compounds with well-studied DNA precipitating properties (Ouameur and Tajmir-Riahi 2004; Deng and Bloomfield 1999) and selected spermine and hexamminecobalt(III) chloride (CoHex) for testing DNA precipitation and recovery profiles using glass beads (Supplemental Figure S4a). Optimum final concentrations of the compounds were tested empirically based on previous studies (Pelta 1996).

The ULK001 protocol is carried out in a dilute condition of ∼50 ng/μl UHMW DNA. We tested the impact of DNA concentration (10, 20 and 50 ng/μl) on precipitation and recovery rate. CoHex is the best precipitant for both recovery and homogenisation of UHMW DNA (Supplemental Figure S4b-d). The optimal DNA recovery of ∼75% was obtained using CoHex at DNA concentrations of 20 ng/μl. Spermine shows a higher recovery rate at 50 ng/μl DNA but greater heterogeneity and sample-to-sample variation (Supplemental Figure S4b-d).

The early ULK001 kit version used a spermine-based precipitation buffer (PPT buffer). We tested this alongside our *FindingNemo* and spermine clean-up protocols, using the same adapted DNA input. These runs yielded similar UL sequencing metrics (Figure 3a). Albeit with a slightly longer N50, ONT’s PPT protocol showed the lowest yield per pore, presumably from the lack of any washing step after the library precipitation that might cause the library to have a higher rate of blocking than the others (Figure 3b).

**Figure 3.**
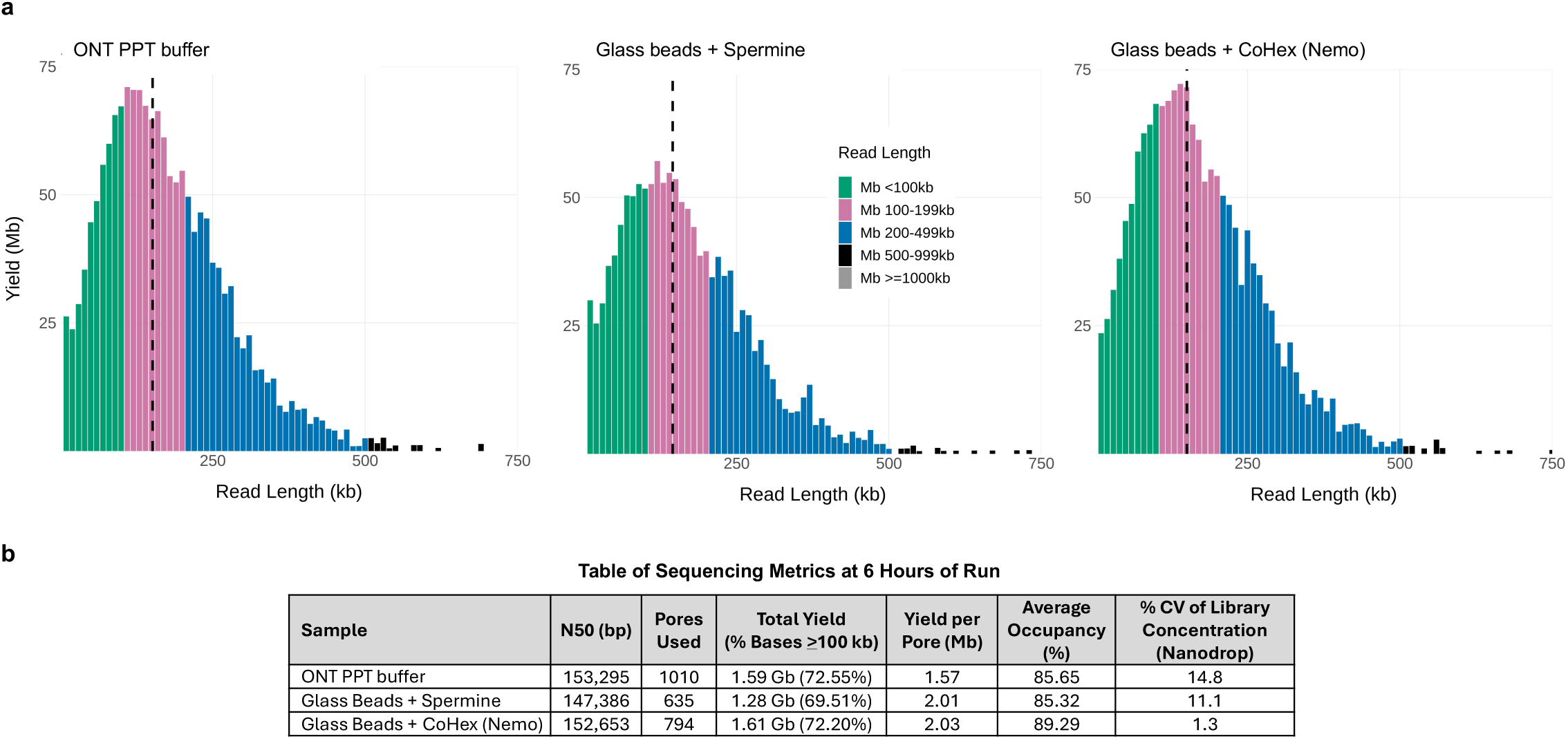
Choosing the best precipitating agent that ensures fast homogenisation: (a) Read length distribution of libraries precipitated with different buffers; dashed black vertical lines denote N50s, (b) Sequencing metrics after 6 hours of run (excluding the first 10 minutes). All libraries were extracted from GM12878 cells using Monarch direct lysis protocol, loaded on MinION R.9.4.1 flow cells and sequenced on GridION platform.

Homogeneity is measured by calculating the coefficient of variation percentage (%CV) of the DNA concentration. We take at least three concentration measurements from the top, middle and bottom part of the solution using a Nanodrop^TM^ spectrophotometer device. The smaller the %CV, the more homogeneous, and vice versa. Recovered DNA became more homogeneous after longer in solution, except when precipitated using CoHex at 10 ng/μl DNA concentration (Supplemental Figure S4e). Previous studies showed that a charge neutralisation of 88-90% is required for DNA condensation (Deng and Bloomfield 1999). We hypothesise that at 10 ng/μl DNA, excess CoHex cations affected the water hydration at the DNA-cations interface (Deng and Bloomfield 1999) resulting in a more condensed, difficult-to-dissolve DNA as observed during pipetting. Moreover, Nanodrop is more sensitive when measuring absorbance of the differentially condensed DNA in the solution which explains the larger variation in DNA concentration compared to Qubit^TM^ measurements (Supplemental Figure S4c-d). In contrast, spermine-precipitated DNA shows the least homogeneity in solution (Supplemental Figure S4e). Spermine is also less stable than CoHex; as a solution it is readily oxidised at room temperature (sigmaaldrich.com). We also tested spermidine as a commonly used DNA precipitation agent but found it less reliable for the recovery of input DNA (data not shown). CoHex was chosen to precipitate DNA at a concentration range of 20-40 ng/μl UHMW DNA.

The importance of DNA homogeneity and optimum concentration during UL library preparation was shown in the use of a concentrated and non-homogeneous UHMW DNA in sequencing, i.e. more than 60 ng/μl DNA in reaction (Supplemental Figure S5). The sequencing output of this library typifies the output of an undercut library where N50 reaches above 100 kb, but accumulated yield and occupancy are low, as well as a rapid decrease of sequencing pore number (Supplemental Figure S5d). In essence, ensuring the homogeneity of extracted UHMW DNA and setting the reaction concentration below 50 ng/μl provide the optimum conditions for UL sequencing. Longer elution of the library (e.g. overnight) at room temperature with gentle rotation may also help untangle long DNA molecules, facilitating further homogenisation and eventually improving UL sequencing output.

#### Quantification of UHMW DNA

One of the complications in UL sequencing is in accurately quantitating the UHMW DNA. There is often large variation between the two most used quantification methods: the spectrophotometric and fluorometric based methods, represented by Nanodrop^TM^ and Qubit^TM^ 3 respectively (Supplemental Figure S4f). The viscous nature of UHMW DNA also makes it difficult to withdraw a homogeneous small volume to be measured, even when using a positive displacement pipette.

Therefore, we followed a novel Qubit concentration measurement of UHMW DNA to complement the standard Nanodrop method. This method uses Jurkat genomic DNA as the baseline standard for UHMW DNA and a glass bead to homogenise samples before each measurement (Koetsier 2021). We modified it by combining small volumes from 3-4 different parts of the DNA solution to average out concentration differences within the solution (Cahyani 2021a). An accurate concentration measurement requires an ideal ratio of Nanodrop to Qubit DNA values between 1-1.5 (Simbolo et al. 2013).

As expected, sheared DNA homogenised faster and better than UHMW DNA as shown by the smaller %CV of the sample a few hours after elution (Supplemental Figure S6a). This fits with the fact that shorter DNA molecules will diffuse faster than longer ones. Longer DNA molecules may also have more complex structure and behaviour in solution (Robertson and Smith 2007). The %CV of both types of samples were all below 50% indicating that the *FindingNemo* method could homogenise DNA solution relatively quickly (Supplemental Figure S6a). It also filtered out contaminants in the input DNA as shown by the increase of the Nanodrop absorbance values after purification (Supplemental Figure S6b) (Koetsier and Cantor 2019). Lastly, the *FindingNemo* protocol did not interfere with the DNA quality as shown by their size distribution on the pulsed-field electrophoresis gel before and after DNA clean-up (Supplemental Figure S6c).

#### The *FindingNemo* protocol vs undercut UL libraries

Combining the *FindingNemo* approach with either the Quick’s or KSF protocols improved sequencing yield but, surprisingly, did not significantly improve occupancy or N50 (Supplemental Table S3). We hypothesise that the transposase reaction upstream of the library clean-up is a more critical step in producing optimal ultra-long sequencing outputs. Suboptimal transposase fragmentation may result in long DNA molecules without adapters in the library, due to an insufficient number of transposome complexes compared to DNA molecules, a situation we refer to as an ‘undercut.’

An undercut library may also occur if the transposome complex-to-DNA ratio is balanced, but the DNA solution is not homogeneous. Viscous, non-homogeneous UHMW DNA is difficult to dilute and can have varying DNA densities at different points in the solution. Consequently, the cutting frequency of the transposome complex will be higher where DNA is more accessible and lower where long DNA molecules remain in a compacted, less accessible mass. This scenario counterintuitively results in a library with a short N50 despite the high molecular weight nature of the input DNA (Supplemental Figure S8 - 0 rpm sample).

It is also important to consider the interaction between UHMW DNA molecules and the sequencing pores, especially just before a mux scan occurs. During this period, only pores with actively threading (sequencing) DNA molecules will continue sequencing; pores without DNA are inactivated. This process terminates at a fixed time, which is 10 minutes by default. After this period, all pores are inactivated. Molecules still traversing a pore at that point will be ejected, but UHMW DNA increases the risk of stalling or blocking nanopores. At a sequencing speed of 400 bases per second, any molecule longer than 240 kb is at risk of blocking a pore and negatively impacting yield. Therefore, in the *FindingNemo* protocol, we modified this script parameter (“wind_down”) from 600 to 1800 seconds (Cahyani et al. 2024).

### C. The *FindingNemo* protocol in UHMW DNA extraction

#### Extractions without kits

We next compared the effect of DNA extraction methods on UL sequencing. The original Quick et al. and Logsdon et al. approaches utilised phenol-chloroform isolation and alcohol-based precipitation with some differences (Table 2A,B). We succeeded in scaling-down these protocols in combination with glass beads to use less cell input and shorten hands-on time to at least half (Table 2C). To reduce the toxicity from phenol use, we also developed phenol-free glass beads-based extraction protocols utilising either SDS, CTAB, or CTAB with CoHex as the lysis buffer (Table 2D, E). These extraction protocols produced (U)HMW DNA, characterised by the distinctly narrow vertical lines/smears and DNA in the wells of the pulsed-field gel electrophoresis results (Supplemental Figure S7).

**Table 2.** Compilation of UHMW DNA extraction methods for UL sequencing, optimised in human GM12878 cells.

|  | Without Kits (Phenol-based) |  |  | Without Kits (Phenol-free) |  | Kit-based |  |
| --- | --- | --- | --- | --- | --- | --- | --- |
|  | Quick's Phenol Protocol (Quick, 2018) | Logsdon's Phenol Protocol (Logsdon, 2020) | Phenol + Glass beads | CTAB or SDS Lysis + glass beads | CTAB-CoHex Lysis + glass beads | NEB Monarch Cell & Blood Kit | Circulomics Nanobind CBB Big DNA Kit |
|  | A | B | C | D | E | F | G |
| Standard cell input | 50 million | 20-70 million | 1-5 million | 1-5 million | 1-5 million | 0.5-5 million | 6 million |
| Phenol-chloroform | Yes | Yes | Yes | None | None | None | None |
| Lysis buffer | SDS | SDS | SDS | CTAB or SDS | CTAB and CoHex | Proprietary | Proprietary |
| SPRI Substrate | None | None | 3-mm glass beads | 3-mm glass beads | 3-mm glass beads | 4-mm glass beads | Nanobind disc |
| Precipitation reagents | Ethanol, Ammonium Acetate | Ethanol, Ammonium Acetate | Ethanol, Ammonium Acetate | Isopropanol, Ammonium Acetate | CoHex | Isopropanol, Proprietary binding buffer | Isopropanol, Proprietary binding buffer |
| DNA extraction time | 4 hours | 4 hours + overnight | 2 hours | 1 hour | 1 hour | ~30 min | 2 hours |
| DNA resuspension time | 2 days | 2 days | min. overnight | min. 2 hours | min. overnight | min. 1 hour (standard overnight) | overnight |

We also successfully produced UL runs using these DNA samples with varying degrees of occupancies and yields (Figure 4). Only the extraction protocol using a combination of CTAB and CoHex in the lysis buffer could not produce UL N50 (Figure 4f). The inclusion of CoHex during lysis was to test if stable DNA toroid aggregates could be induced (Deng and Bloomfield 1999; Ouameur 2004), so that DNA length could be kept intact during extraction and eventually improve sequencing output. The CoHex precipitated DNA was then washed with ethanol, and it rendered DNA elution from the beads more difficult. Ethanol is required in the wash buffer as it rinses salt, contaminants and other impurities from the cell extract while simultaneously affecting DNA condensation (Arscott et al. 1995; Oda et al. 2016). More optimisation is needed to obtain a consistent rate of elution and recovery in the CTAB-CoHex lysis protocol. Nevertheless, the option of using one of two oppositely charged surfactants as lysis agents, i.e. cationic CTAB (Arseneau et al. 2016) and anionic SDS (Xia et al. 2019), provides flexibility in extracting UHMW DNA from diverse sample types and context.

**Figure 4.**
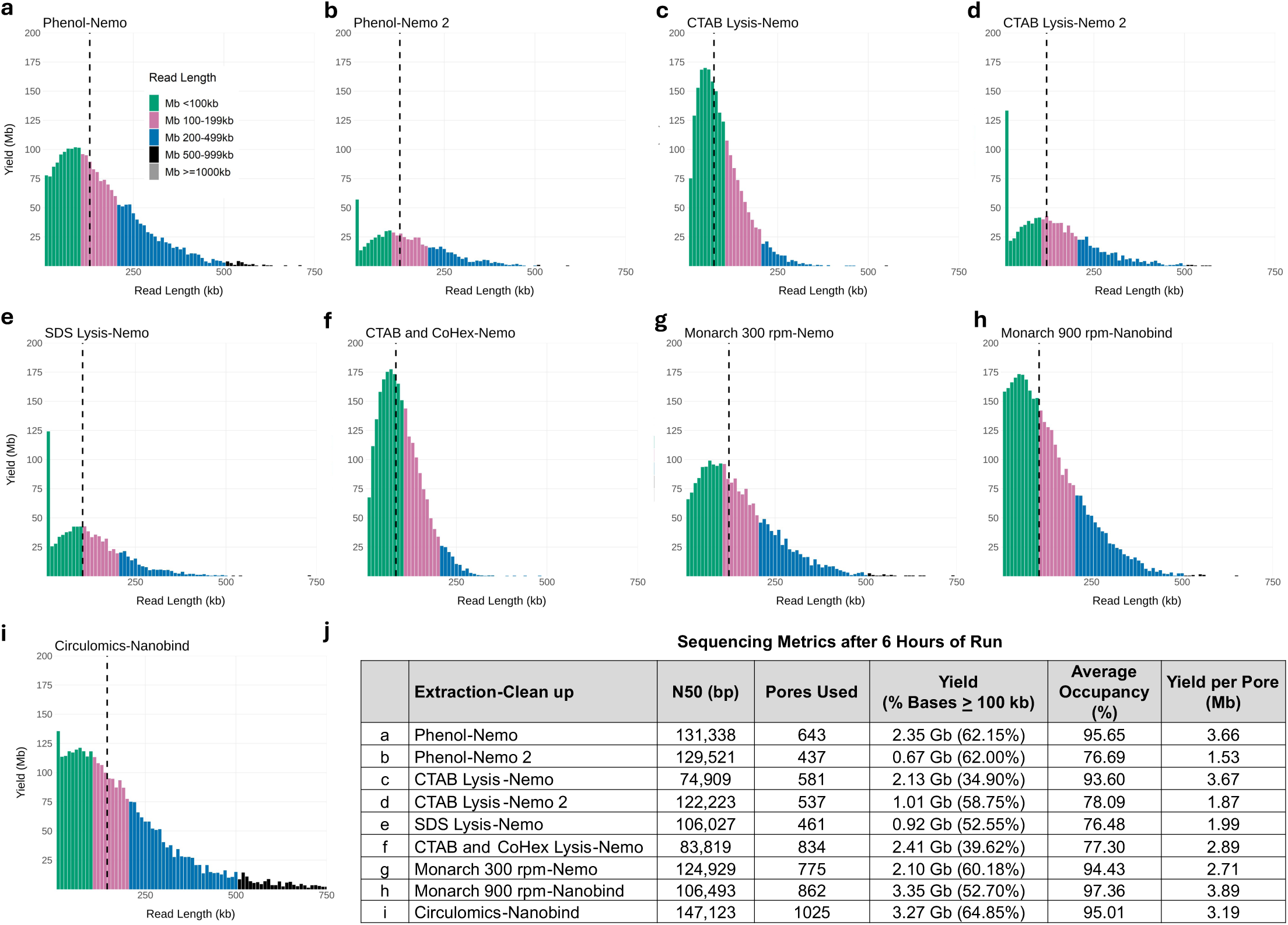
Combinatorial effects of extraction and library clean-up methods, either with or without the use of kits, on the UL sequencing outputs: (a-i) Read length distributions of the libraries: graph titles denote the used extraction kit/method and the clean-up protocol, dashed black vertical lines denote N50s. (j) Sequencing metrics after 6 hours of run (excluding the first 10 minutes). DNA was extracted from GM12878 cells. Each library was loaded on a MinION R9.4.1 flow cell and run on the GridION platform.

#### Extraction using kits

We tested a range of (U)HMW DNA extraction kits during our method development and found that Nanobind CBB Big DNA (Circulomics) and Monarch (NEB) were best suited for our purposes (Table 2F-G). The Nanobind CBB Big DNA Circulomics kit could be used to extract HMW or UHMW DNA, depending on the chosen protocol. DNA extracted with the UHMW protocol showed higher size distribution on a pulsed-field electrophoresis gel compared to the HMW protocol (Supplemental Figure S7a vs S7b). Libraries prepared from UHMW DNA samples extracted using the Circulomics kit produced UL reads and good sequencing metrics (Figure 4i-j).

The Monarch kit protocol initially involved a two-step DNA extraction: first extracting nuclei from cells, then isolating DNA from the nuclei (“Nuclei Prep”). It was later refined to a “Direct Lysis” method, which bypassed the “Nuclei Prep” step by lysing cells and nuclei simultaneously to release DNA. This method yielded high-quality UHMW DNA from GM12878 cells, as shown in pulsed-field gel samples (Supplemental Figure S7a, b). Libraries prepared from these DNA samples indicated that the Direct Lysis approach performed slightly better than the Nuclei Prep method (Supplemental Figure S8).

We also found that the shaking speed during lysis in the Monarch protocol influenced DNA homogeneity, which affected sequencing N50 and occupancy (Supplemental Figure S8; Figure 4g-h; Supplemental Table S4). Lower lysis speeds yielded longer DNA fragments by keeping DNA more intact. However, no shaking during lysis could result in a non-homogeneous UHMW DNA solution, negatively affecting the transposase reaction during library preparation (Supplemental Figure S8 - 0 rpm). This resulted in a library with non-UL read N50, the lowest occupancy, and yield compared to other libraries (Supplemental Figure S8). The best UL output was achieved by preparing libraries from DNA lysed at a medium speed of 600 to 900 rpm (Supplemental Figure S8; Figure 4g-h; Supplemental Table S4).

### D. Sequencing parameters and performance of the *FindingNemo* protocol

#### Maximum yield is obtained at a read N50 between 90-110 kb

On a MinION, yields in excess of 3 Mb per pore after 6 hours of sequencing could be obtained at 93% occupancy, reinforcing that high occupancy is essential for high yield (Figure 4j). However, UL libraries with many short fragments had reduced occupancy as short reads pass quickly through the pores, lowering total yields (Figure 4b, d, e, j). This underscores the importance of homogeneously cut UHMW DNA in UL library preparation. Additionally, targeting an N50 between 90-110 kb is ideal for optimal ultra-long sequencing output, as N50 is inversely proportional to yield (Figure 4j; Supplemental Table S4)

#### Scalable library loading amount with the *FindingNemo* protocol

We also tested the impact of the amount of input DNA and library loaded per flow cell. When the input amount of UHMW DNA was less than 5 μg, sequencing output parameters were markedly reduced (Supplemental Table S4, Monarch-01 vs Monarch-05). To produce the optimum N50, occupancy and yield, DNA equivalent to at least one million GM12878 cells (or at least 5 μg) should be used for library preparation, and a minimum of 2 μg of this library loaded onto a MinION flow cell. At least two million cells and 4 μg of library should be used on a PromethION flow cell (Supplemental Table S4, Monarch-01 vs Monarch-02).

#### *FindingNemo* in one day: from cells to UL sequencing in one working day

The relatively fast homogenization rate of the *FindingNemo* clean-up method, combined with the fast DNA extraction when using the Monarch kit can support the preparation of UL libraries from fresh or frozen cells to sequencing in just a working day (Supplemental Figure S9a) (Cahyani 2021b). We can obtain UL N50s using this protocol, however yields are markedly reduced (Supplemental Figure S9b-d), likely as a result of the heterogeneity of DNA samples. Yield reductions were exacerbated by the low occupancy level and the abundance of short reads and would be interpreted as undercut libraries (Supplemental Figure S9b-d; Run 2 and 3). Moreover, the *FindingNemo* clean-up protocol might not significantly remove these highly abundant short reads due to their high concentration in the libraries.

### E. Using the *FindingNemo* protocol in diverse sample types, chemistry and platforms

#### The *FindingNemo* protocol works with the ULK114 new chemistry kit

In 2022, ONT released a new pore and chemistry running on the R10.4.1 flow cells as well as a new ultra-long kit, ULK114. This included new sequencing scripts and a sample rate shift from 4 kHz to 5 kHz. We tested the ULK114 kit according to the manufacturer’s instructions and with the *FindingNemo* protocol on two human cell lines (Figure 5). Libraries prepared with the *FindingNemo* protocol showed longer N50s, optimal yield, and occupancy (Figure 5a-b). This protocol consistently produced higher yields, regardless of kit chemistry and sequencing programs. Additionally, the 5 kHz script increased the overall mapping rate of reads compared to the 4 kHz program (Figure 5b).

**Figure 5.**
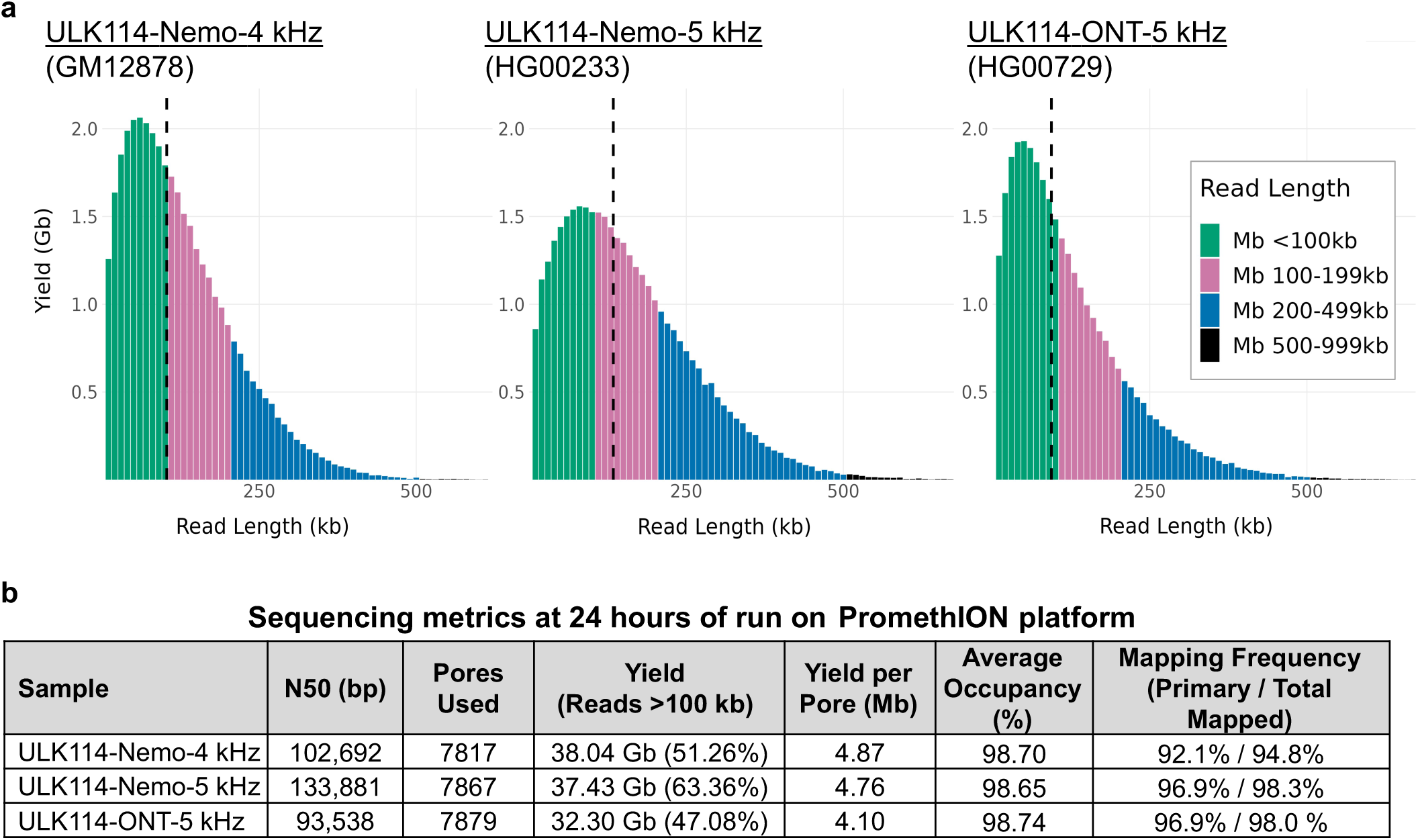
Ultra-long N50s were obtained using the new ULK114 kit from three cell lines: (a) Read length distribution of GM12878 library run with the earlier ULK114 script version (left panel), while the other two libraries (middle and right panel) were run with the most current 5 kHz version of the script. Vertical dashed black lines indicate N50s. (b) Run metrics of the three libraries showing the Nemo protocol performed better than ONT’s.

#### Ratio of transposase to DNA

The ratio of transposase to DNA is crucial for optimising UL sequencing. The ULK001 protocol specifies 1 μl of transposase fragmentation mix (FRA) per 1 million human cells, corresponding to 5-6 μg of UHMW DNA for a 3.2 Gb genome (Table 3). We used this ratio for all GM12878 (Figures 2-5) and HEK293 (Figure 6) runs. Additionally, we tested different FRA to DNA ratios in Mongolian gerbil, tick *Amblyomma variegatum*, and yeast *Saccharomyces cerevisiae* (Table 3). These ratios produced UL sequencing outputs for the gerbil and tick samples (Figure 6a, c). The read N50 for the tick exceeded its genome assembly’s contig N50, indicating significant assembly improvement (Alistair Darby, personal communication). Although the yeast run did not yield UL read N50s (Figure 6b, c), it showed a significant increase in read length compared to existing nanopore-sequenced data (Giordano et al. 2017; Salazar et al. 2017).

**Figure 6.**
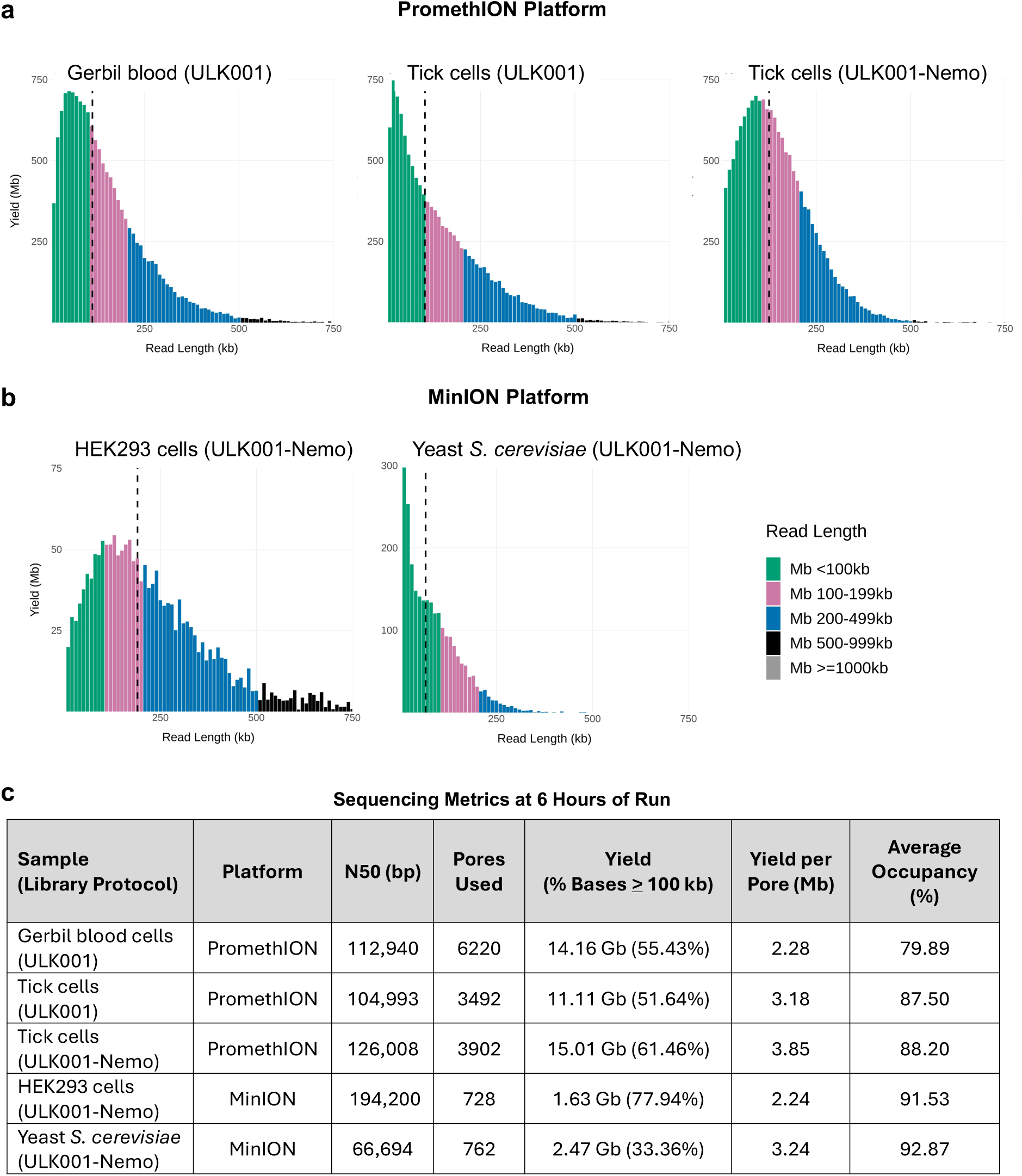
Ultra-long sequencing output and metrics of non-GM12878 cells and different sequencing platforms: gerbil blood, tick *Amblyomma variegatum*, yeast *Saccharomyces cerevisiae* and human HEK293 cells using either the ULK001 or the ULK001-Nemo protocol (as labelled). (a) Read length distributions of libraries run on PromethION R9.4.1 flow cells; dashed black vertical lines denote N50s, (b) Read length distribution of the HEK293 and yeast *S. cerevisiae* libraries run on MinION R9.4.1 flow cells, (c) Sequencing metrics of the libraries. Each metric was shown after 6 hours of run (excluding the first 10 minutes).

**Table 3.** Ratio of FRA to DNA in different genomes.

| Species | Genome Size | Cell Input Number | DNA Equivalent (µg) | FRA Volume (µl) |
| --- | --- | --- | --- | --- |
| Human GM12878 | 3.2 Gb | 1 x 10<sup>6</sup> | 5-6 | 1 |
| Human HEK293 | 3.2 Gb | 1 x 10<sup>6</sup> | 6-8* | 1 |
| Mongolian Gerbil | ~2.7 Gb | Unknown | 10 | 1.5 |
| Tick *A. variegatum* | ~6.0 Gb | 1 x 10<sup>6</sup> | 11-13 | 2.5 |
| Yeast *S. cerevisiae* (n) | 12 Mb | 2 x 10<sup>8</sup> | 4-5 | 6 |

(\*) extracted DNA yield is likely to be more than normal diploid human cell line because of its hypotriploid nature (Lin 2014)

#### Genome ploidy

Genome ploidy anomalies must be considered when preparing UL libraries. For instance, about 4.2% of HEK293 cells are hypotriploid (Synthego HEK293). This means the DNA mass extracted from the same number of cells is higher than from the GM12878 cell line, altering the FRA to DNA ratio (Table 3) and potentially affecting sequencing output by increasing read length and decreasing yield. Sequencing HEK293 DNA with the Monarch kit confirmed this, producing N50s over 190 kb but with lower yields (Figure 6b, c). To maximise yield and N50, it is better to use the FRA volume to DNA mass ratio rather than the absolute cell number when genome ploidy information is available.

#### Sample types: to count or to weigh?

The UL protocol requires the same number of human cells per microliter of FRA, whether from a cell line or nucleated blood. However, counting cells in blood samples can be impractical, as seen in the gerbil UL run (Table 3). In such cases, the FRA to DNA mass ratio can be used, provided the DNA is extracted to maintain native chromosome length and properly quantified (Methods). Using this ratio, UL libraries were produced from gerbil blood following the original ULK001 protocol (Figure 6a, c).

#### The *FindingNemo* protocol works in different nanopore platforms

We ran pairwise sequencing of identical libraries on both the MinION and PromethION platforms, showing that PromethION produced more yield per pore (Supplemental Table S5). With nearly six times as many pores as a MinION, PromethION is expected to generate more data. The best PromethION results had an N50 of 126 kb and a yield of 3.85 Mb per pore at 88% occupancy (Figure 6c - Tick cells ULK001-Nemo). On the MinION platform, the best results ranged from 106-147 kb N50, with yields of 3.19-3.89 Mb per pore and at least 95% occupancy (Figure 4j). These metrics were all obtained after 6 hours of sequencing.

#### Relative performance of extraction kits in tick samples

We compared two extraction kits, Circulomics and Monarch, for UL sequencing of a tick cell line with a ∼6 Gb genome (Figure 6a, c). Although flow cell occupancies were similar, the Monarch-extracted DNA had longer N50s, higher yields, and a greater proportion of UL reads (Figure 6a right panel, c). The Circulomics library had more short reads, likely due to undercutting (Figure 6a middle panel). The Circulomics-extracted DNA concentration was ∼50 ng/μl, while Monarch-extracted DNA was ∼26 ng/μl, reflecting differences in sample homogeneity. Moreover, Monarch extractions yielded DNA with a longer size distribution, as shown by pulsed-field electrophoresis gel (Supplemental Figure S10). The Monarch protocol is also more flexible, allowing modifications to target different read-length distributions by adjusting lysis speed (Methods).

## Discussion

High-quality, uniformly long DNA is essential for ultra-long reads on Oxford Nanopore sequencers. Our results support the hypothesis that to obtain this length uniformity, transposase reactions should occur in a dilute and homogeneous DNA solution. This can be achieved in a large reaction volume and by thorough mixing to obtain a ‘properly cut’ library in contrast to ‘undercut’ as explained before, or ‘overcut’ which is a higher ratio of FRA to DNA molecules (Figure 7). Additionally, non-adapted DNA molecules may ‘crowd’ the space around the pores thereby decreasing the diffusion rate of the adapted DNA molecules to be tethered to the pores. The adapted long DNA molecules themselves may inactivate or block the pores during sequencing because of their lengths.

**Figure 7.**
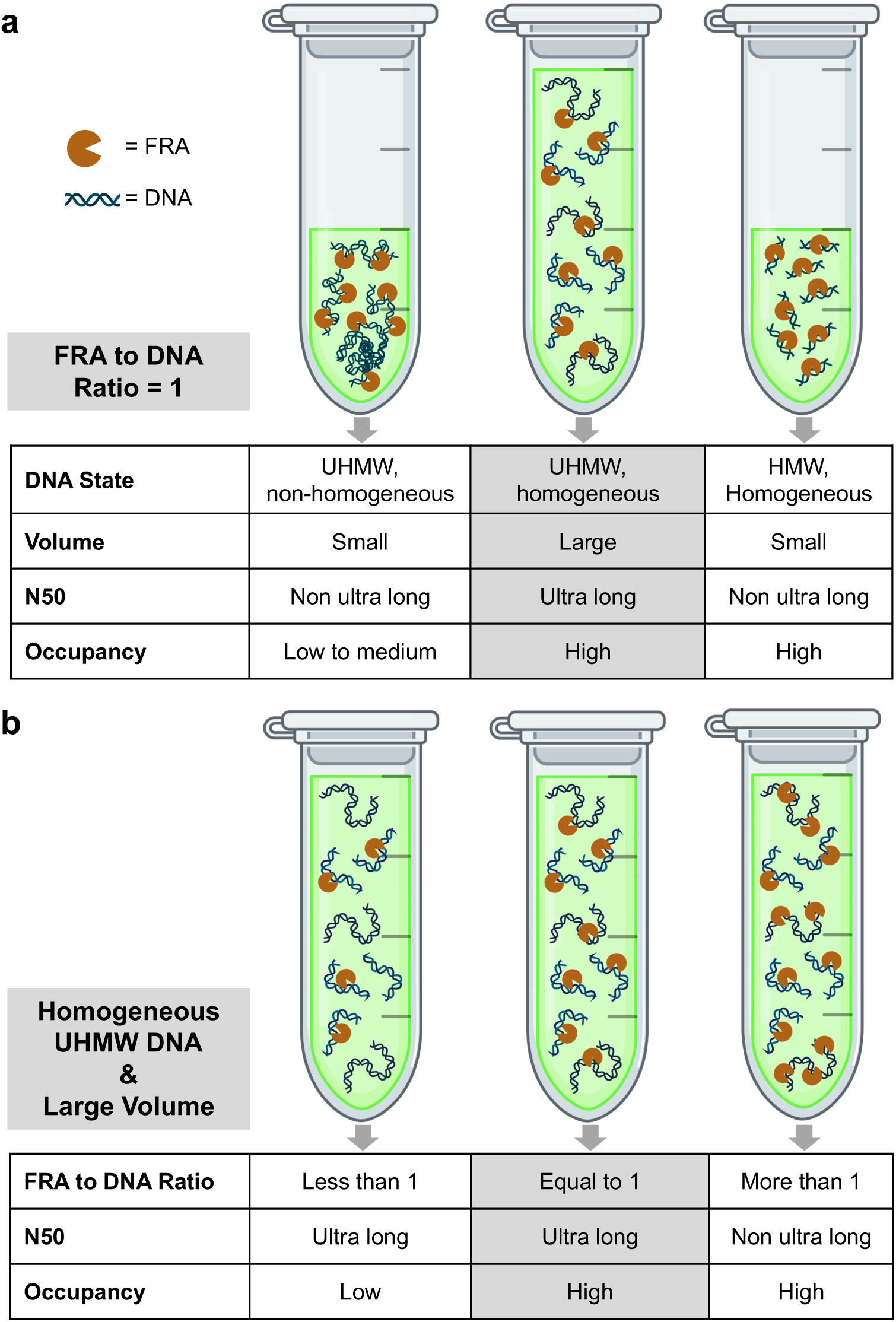
Optimum conditions for ultra-long sequencing - to obtain an optimum UL sequencing output, there needs to be a 1:1 ratio of FRA to DNA molecules in a homogeneous solution ascertained by proper cell counting and/or DNA mass measurement. [Image created with BioRender.com]

With these challenges, control of DNA quality and quantity is of primary importance. Proper quantification of the number of input cells or tissue weight has direct consequence to controlling DNA yield and to some extent, length distribution. Extended elution and homogenisation are also beneficial; the longer that an UHMW DNA sample is allowed to equilibrate, the more homogeneous it becomes. This reduces the benefit of our *FindingNemo in One Day* protocol where both maximal read lengths and yields are important. However, as we show, a rapid UL library preparation is possible, albeit with lower total yield. Further optimisation is required, especially at the homogenization of DNA post-extraction, to make this protocol more robust.

Chromosome length distribution of a genome may limit the maximum read N50 that can be obtained, as was the case with the yeast *S. cerevisiae* ultra-long library. Prior knowledge of the genome size and chromosome lengths combined with empirical tests may be required to find the optimum ratio of transposase to DNA amount and produce maximum sequencing N50 and yield. This is a challenging part of ultra-long sequencing as it needs fine tuning and adjustments depending on the sample types used.

We anticipate that the *FindingNemo* protocol will be generally applicable to tissue samples, could be readily multiplexed and is likely compatible with automation. We have also applied the *FindingNemo* clean-up to ONT’s ligation-based protocols, e.g. SQK-LSK114, and can obtain UL libraries although results were not consistent in both read length and yield (data not shown). We suspect there may be incompatibility in the underlying chemical properties and reactions behind (U)HMW DNA, ligation buffer compounds and CoHex that made the final DNA elution difficult. As the components of the ligation buffer are proprietary to ONT, more empirical optimization is required for this protocol. Using the *FindingNemo* protocol in a ligation-based library preparation may potentially and significantly reduce the required input DNA amount and enable processing of unsheared HMW DNA, which cannot be cleaned by the standard magnetic SPRI-based methods. If successful, this can also be applied to the recently developed ONT duplex sequencing and thus produce an ultra-long duplex sequencing approach.

Finally, open-source development and sharing of protocols is crucial to enable the widest access to cost-effective high throughput ultra long sequencing and troubleshooting. We established the LongRead Club to house these protocols and promote reproducibility (LongRead Club, n.d.). The *FindingNemo* protocol described here is a valuable addition to the repertoire of ultra-long nanopore sequencing methods available.

## Methods

### A. Genomic DNA extraction from cells

#### Cell sources

GM12878 cells were grown by Darren Crowley (University of Nottingham) and additionally purchased from Coriell Institute for Medical Research, USA. Yeast *S. cerevisiae* cells were given by Stephen Gray (University of Nottingham). Tick *A. variegatum* (AVL/CTVM17) cells were obtained from Alistair Darby’s group (University of Liverpool). The HG00233 and HG04054 cells were obtained from Danny Miller’s lab (University of Washington, USA). Lastly, HEK293 cells were obtained from New England Biolab (NEB) in Ipswich, MA, USA.

#### Kit-based extraction

Kits used in the extraction of UHMW DNA were Monarch® HMW DNA Extraction Kit for Cells & Blood (NEB T3050) and Nanobind CBB Big DNA Kit (Circulomics SKU NB-900-001-01) plus the auxiliary Nanobind UL Library Prep Kit (Circulomics SKU NB-900-601-01) according to the manufacturer’s protocols with modifications as previously described for the Monarch protocol (Cahyani 2021a). The Monarch nuclei prep approach followed the original protocol (NEB T3050) while the direct lysis approach combined the prep and lysis buffers into one step (NEB Direct Lysis). Lysis speed at 600 up to 900 rpm resulted in an optimum UL sequencing output and a minimum of overnight incubation of DNA in solution helped homogenization.

#### Phenol-chloroform extraction with glass beads

This is a scaled-down version of Quick’s original protocol (Quick 2018) and the modifications are as previously described (Cahyani 2021a). In summary, pellet of 5 million cells was washed with PBS and lysed with an SDS buffer [0.5% SDS, 100mM NaCl, 25mM EDTA pH 8.0, 10mM Tris-HCl pH 8.0] in the presence of RNaseA [20 μg/ml] at 37 °C for 5 min. Proteinase K [200 μg/ml] was added and incubated at 56 °C for 15 min. Cell lysate was split into two 5PRIME phase-lock gels (Quanta-Bio 2302820). BioUltra TE-saturated phenol (Merck 77607) was added at 1:1 volume ratio. Sample was homogenised by vertical rotation at 20-30 rpm for 10 min, then centrifuged at 4000 x g for 10 min. The aqueous phase was transferred into new phase-lock gel tubes. DNA purification was repeated with a second round of phenol:chloroform:isoamyl alcohol = 25:24:1 volume ratio. In the third round, only chloroform:isoamyl alcohol=24:1 was used. After the last centrifugation, the aqueous phase from the two tubes was combined in a 5 ml or 15 ml tube and added with 5M ammonium acetate (0.4 volume), 3 glass beads (3-mm diameter), and absolute ethanol (2.5 volume). The tube was inverted by hand 20-30 times to precipitate DNA onto the glass beads (or placed in a tube rotator at 10 rpm for 3 min). The supernatant was removed, and bound DNA was washed twice with 60-70% ethanol. Beads with DNA were poured into a bead retainer and quickly spun to remove excess ethanol (or absorbed with a filter paper or tissue). DNA was quickly eluted from beads by pouring them into a 2 ml LoBind tube containing 10mM Tris-HCl pH 9.0 and incubating at 37 °C for 30 min with regular wide-bore pipette mixing. Incubation was continued overnight at room temperature. Afterwards, using a bead retainer, DNA was spun down at maximum speed for 1 min.

#### Phenol-free, home brew extractions

This protocol is adapted from the Monarch® HMW DNA Extraction Kit for Cells & Blood as described previously (Cahyani, 2021-1). In summary, a pellet of 1-3 million cells was washed with PBS and lysed in either SDS or CTAB buffer with 200 μg Proteinase K and incubated at 56 °C for 15 min (preferably with shaking at 600-700 rpm). The SDS buffer composition was the same as used in the phenol-based extraction described in the previous section. The CTAB buffer compositions were 2% CTAB, 1.4M NaCl, 25mM EDTA pH 8.0, and 10mM Tris-HCl pH 8.0. After lysis, 100 μg RNaseA was added and incubated at 37 °C for 5 min. To precipitate DNA, 5M ammonium acetate was added (0.4 volume), 2-3 glass beads (depending on cell number used), and isopropanol (0.9 volume). Sample was mixed on a rotator at 9 rpm for 5 min (or inverted by hand 20-30 times). Liquid was removed by pipetting and bound DNA was washed twice with 60-70% ethanol. Beads with DNA were poured into a bead retainer and quickly spun to remove excess ethanol (or absorbed with a filter paper or tissue). DNA was quickly eluted with 10mM Tris-HCl pH 8.0 in a 2 ml tube and incubated at 37 °C for 30 min with regular wide-bore pipette mixing. Incubation was continued overnight at room temperature. Afterwards, using a bead retainer, DNA was spun down at maximum speed for 1 min.

#### CTAB-CoHex extraction

This protocol is similar to the home-brew CTAB method described above with the following modifications. The CTAB-CoHex buffer compositions used were 2% CTAB, 20mM CoHex, 1.4M NaCl, 25mM EDTA pH 8.0, and 10mM Tris-HCl pH 8.0. To precipitate DNA, sample was diluted with 10mM Tris-HCl pH 8.0, so CoHex concentration became ∼5mM and NaCl was ∼400mM, *i.e*. ratio of CTAB-CoHex lysis buffer to Tris-HCl was 1:2.5. Then, 2-3 glass beads (depending on cell number used) were added. Sample was mixed on a rotator at 9 rpm for 5 min (or inverted by hand 20-30 times). Liquid was removed by pipetting and bound DNA was washed twice with an ethanol buffer (50% ethanol, 50% of 1mM CoHex). Beads with DNA were poured into a bead retainer and quickly spun to remove excess ethanol (or absorbed with a filter paper or tissue). DNA was quickly eluted with 10mM Tris-HCl pH 8.0 in a 2 ml tube and incubated at 37 °C for 30 min with regular wide-bore pipette mixing. Incubation was continued overnight at room temperature. Afterwards, using a bead retainer, DNA was spun down at maximum speed for 1 min.

#### Yeast extraction using CTAB and glass beads (Cahyani 2024)

Yeast pellet (∼50 μl volume or equivalent to 200 million cells) was washed twice in cold PBS and resuspended in 480 μl SpheroBuffer (1M Sorbitol, 100mM Na_2_HPO_4_, 100mM EDTA). Lyticase enzyme (5U/μl) was added (20 ul). Sample was incubated at 30°C for 30 minutes and spheroplast formation was checked from time to time. Sample was lysed by adding 480 μl of 2X CTAB lysis buffer (4% CTAB, 2M NaCl, 50 mM EDTA pH 8.0, and 20 mM Tris-HCl pH 9.0), mixed well with wide-bore pipet tip as the buffer is viscous, added with 400 μg Proteinase K (Qiagen 20 mg/ml) and incubated at 56°C for 15 minutes whilst shaking at 700 rpm. To remove RNA, 400 μg of RNaseA (Qiagen 100 mg/ml) was added followed by incubation at 56°C for 10 minutes without shaking. Sample was cooled off at room temperature and added with 400 μl of 5M ammonium acetate and 100 μl of 5M NaCl and mixed by a vertical rotator for 1 minute at 9 rpm. Sample was then centrifuged for 7 minutes at 16,000x g and the lysate supernatant was removed to a new 2 ml tube. Three glass beads and 0.5 ml isopropanol were added, and the sample was mixed on a vertical rotating mixer at 9 rpm for 5 minutes. Between 0.5-0.7 ml liquid was removed and replaced with the same amount of isopropanol and was rotated for an additional 3 minutes. When DNA had bound tightly around the beads, the liquid was discarded, and bound DNA was washed twice with 1 ml of 70% ethanol. Beads with DNA were poured into a bead retainer and quickly spun to remove excess ethanol (or absorbed with a filter paper or tissue). DNA was quickly eluted with EB (ONT) or 10mM Tris-HCl pH 9.0 in a 2 ml tube and incubated at 37 °C for 30 min with regular wide-bore pipette mixing. Incubation was continued overnight at room temperature. Afterwards, using a bead retainer, DNA was separated from the beads by centrifugation at maximum speed for 1 minute.

### B. Quantification of UHMW DNA

Two nucleic acid quantification methods: fluorometric-based Qubit [Thermo Fisher Scientific, MA, USA] and spectrophotometric-based Nanodrop [Thermo Fisher Scientific], were used in parallel to assess both the quantity and purity of the extracted DNA. The quantification follows the published protocol (Koetsier, 2021) with some modification as previously described (Cahyani, 2021-1). In brief, DNA is sampled from four different points in the tube: top, upper-middle, lower-middle, and bottom part of the solution. Then, a glass bead was added, and sample was vortexed at full speed for 1 min for concentration measurements. For Qubit measurement, Jurkat genomic DNA (Thermofisher SD1111) was used as a concentration standard instead of the lambda DNA provided by the manufacturer (‘Giron’ Koetsier and Cantor 2021). To measure DNA homogeneity in solution, %CV of concentration was calculated by directly measuring concentrations of 3-4 different points of the solution using Nanodrop.

#### C. Quality check of UHMW DNA using PFGE

To assess DNA quality and read-length distribution, 200-300 ng extracted DNA was run on 0.75% gold agarose gel (SeaKem 50150). Pulsed-field gel electrophoresis was carried out using Pippin Pulse device (Sage Science) at a pre-set mode of “5-430 kb” for 16 hours. Gel was stained with RedSafe DNA stain (ChemBio 21141) and visualised with a Bio-Rad GelDoc imaging system.

### D. Rapid kit-based UL library preparation

#### Ultra-long kit-Nemo

The ONT ultra-long kits, i.e. ULK001 and ULK114, are based on the cleaving and tagging of the DNA by the transposome complexes to insert transposase adapters, which then are ligated to the sequencing adapters (ONT ULK001). Our modifications of the ULK001 method were described in previously published protocol (Cahyani, 2021-1). To summarise, DNA extracted from 1-6 million cells was diluted to a final concentration of 20-40 ng/μl (in the final reaction volume). All mixing was done with wide-bore pipette tips. Meanwhile on ice, the dilution buffer (FDB) was well mixed with FRA (1 μl per 1 million human cells DNA), then combined to the DNA solution, and quickly incubated at 23 °C for 5-10 min (depending on the reaction volume). Enzyme was inactivated at 70 °C for 5 min and cooled down to room temperature. Sequencing adapter (RAP-F) was added (0.83 μl per 1 μl FRA used) and the sample was incubated at room temperature for at least 30 min.

The ULK114 modifications are similar to ULK001 and are described in the newly published and updated protocol (Cahyani 2024). In summary, library preparation to run on a PromethION flow cell needs extracted DNA from 6 million cells that was diluted to a final concentration of 20-40 ng/μl (in the final reaction volume). On ice, the dilution buffer (FDB) was well mixed with FRA (6 μl per 6 million human cells DNA), then combined to the DNA solution, and quickly incubated at 23 °C for 10 min with 9 rpm rotation. Enzyme was inactivated at 75 °C for 10 min and cooled down to room temperature. Sequencing adapter (RA) was added (5 μl per 6 μl FRA) and the sample was rotated at 9 rpm for 30 minutes.

#### RAD004-Nemo

The RAD004-Nemo protocol followed the same route as the ULK001-Nemo protocol described previously (Cahyani, 2021-1). However, the transposase and adapter used were the ones from the RAD004 kit (*i.e.* FRA and RAP, respectively). The dilution buffer used to replace FDB was the 4X MuA buffer [100 mM Tris-HCl pH 8.0, 40 mM MgCl2, 440 mM NaCl, 0.2% TritonX-100, 40% Glycerol].

#### Krazy StarFish (KSF)

The KSF protocol is an alternative UL sequencing that uses a filter paper instead of glass beads. It is a cheap, quick-and-dirty method to produce UL sequencing based on the RAD004 kit. This method was described previously (Cahyani, 2021-1) and summarised as follows. An amount of ∼7.5 μg DNA in 75 μl volume was mixed well with 25 μl 4X MuA buffer. Meanwhile in another tube on ice, 25 μl 4X MuA buffer was diluted with 74 μl water and added with 1 μl FRA (or up to 1.2 μl FRA if DNA was quite viscous). These two solutions were mixed well on ice, aliquoted into two PCR tubes, and treated at 30° C for 1 min, 80° C for 1 min then cooled to room temperature.

Reactions were pooled into a single tube, 12 μl 5M NaCl added, and mixed gently. A Krazy StarFish filter (made from Whatman filter paper no. 3 with a star hole puncher) was submerged in the solution, 142 μl isopropanol added, and mixed by inversion 20-30 times allowing UHMW DNA to collect and condense onto the filter. Liquid was removed and filter paper was washed twice with 60-70% ethanol. Excess ethanol was removed by pipetting and paper was air-dried for 2 min. After the paper was moved to a clean tube, the library was eluted by adding 120 μl of elution buffer [10mM Tris-HCl pH 8.0]. Before sequencing, 37.5 μl library was added with 0.5 μl RAP and incubated for at least 30 min at room temperature.

### E. Nemo Clean Up: UHMW DNA Library Purification with Glass Beads and CoHex

The Nemo protocol is at the heart of this study and was previously described as an alcohol-free library clean-up method (Cahyani, 2021-1). Typically, for DNA equivalent to 1 million cells or more, 3 glass beads were used. In brief, glass beads were added to the library to be cleaned-up in a 1:1 volume ratio of 10 mM CoHex (2X), i.e. final CoHex concentration is 5 mM. Sample was rotated at 9 rpm for 3 min, or 20-25 times if using hand inversion. Liquid was removed and beads were washed twice with PEGW buffer [10% PEG-8000, 0.5M NaCl] for 3 minutes each. Excess buffer was removed with a fine pipette tip after a quick spin of the tube. Alternatively, the beads with DNA could be poured into a bead retainer (home-made or using the NEB-T3004) and the buffer was absorbed by tapping on a filter paper/tissue. Beads should not be let dry, were quickly eluted with 10mM Tris-HCl pH 9.0 or EB (ONT) and incubated at 37 °C for 30 min with regular wide-bore pipette mixing. Incubation was continued overnight (preferable) at room temperature. Afterwards, using a bead retainer on a new tube, DNA was spun down at maximum speed for 1 min.

### F. Flowcell Priming and Library Loading

All flow cells were primed according to the ULK001 and ULK114 protocols. Library was loaded onto a primed FLO-MIN106 or FLO-PRO002 (R9.4.1), or FLO-MIN114 or FLO-PRO114 (R10.4.1) flow cells for sequencing on a GridION or PromethION, respectively, using the accompanying MinKNOW version software. Loaded UL library was let to tether on the flow cell for at least 30 min before run was started using the ONT ultra-long script with mux scan of every 6 hours for R9.4.1 and the default 1.5 hours for R10.4.1.

### G. *FindingNemo* in one day

To run UL sequencing within the same day as the DNA extraction, *FindingNemo in one day* protocol was developed as previously described (Cahyani, 2021b). In brief, cells were extracted using the Monarch kit and lysed at 700-800 rpm. Eluted DNA was incubated at 37 °C and regularly mixed for at least two hours. After quantification showed that DNA was sufficiently homogeneous (%CV < 100%), the library was prepared following the ULK001-Nemo protocol. Eluted library was further incubated at 37 °C for at least 30 min with regular pipette mixing, quantified and loaded on a primed flow cell.

### I. NanoPlot Data Analysis and Plotting

Violin plots of sequencing speed and distribution graphs comparing read-lengths and read quality were created using the summary file data processed by the NanoPlot tool (de Couster 2023).

### J. R Data Analyses and Plotting

Read-length histograms, occupancy, yield and other sequencing metrics were analysed from the summary, mux, and duty time (or in newer filename version, pore activity) output files using in-house R scripts available on GitHub (https://github.com/cinswasti/FindingNemo).

## Data Access

Most sequencing data generated in this study has been submitted to the European Nucleotide Archive (ENA) under accession number PRJEB76809. Tick sequencing data are currently being analysed as part of an on-going project and will be published in due time. For human work the reference genome used is the hg38 (NCBI RefSeq assembly GCF_000001405.40).

## Acknowledgements

We would like to thank Giron Koetsier (NEB), Simon Mayes (ONT), and Kelvin Liu (Circulomics) for lending their expertise and/or advanced product trials. We thank Darren Crowley and Luke Simpson for supplying the GM12878 cells, Stephen Gray for the yeast cells, Danny Miller and Miranda Galey (University of Washington) for the Human Genome (HG) cells, Lesley Bell-Sakyi and Alistair Darby (University of Liverpool) for the tick *A. variegatum* cells, Giron Koetsier (NEB) for the HEK293 cells, and John Mulley (Bangor University) for the use of the gerbil’s sequencing metrics. John Tyson is grateful to Terrance Snutch. This work was supported by the Wellcome Trust (grant number 204843/Z/16/Z, I.C., N.H., N.L and M.L.).

---

**Figure S1.**
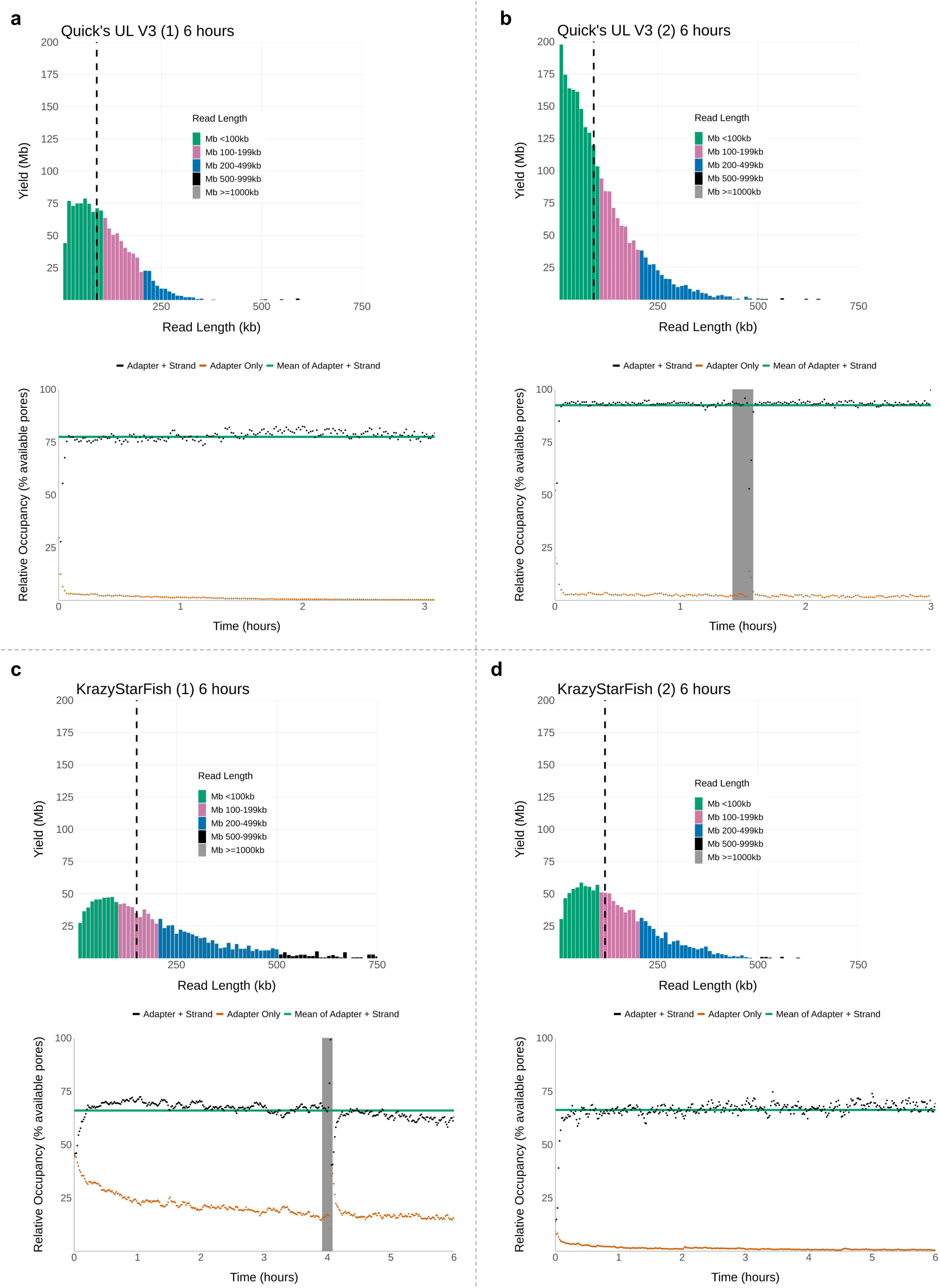
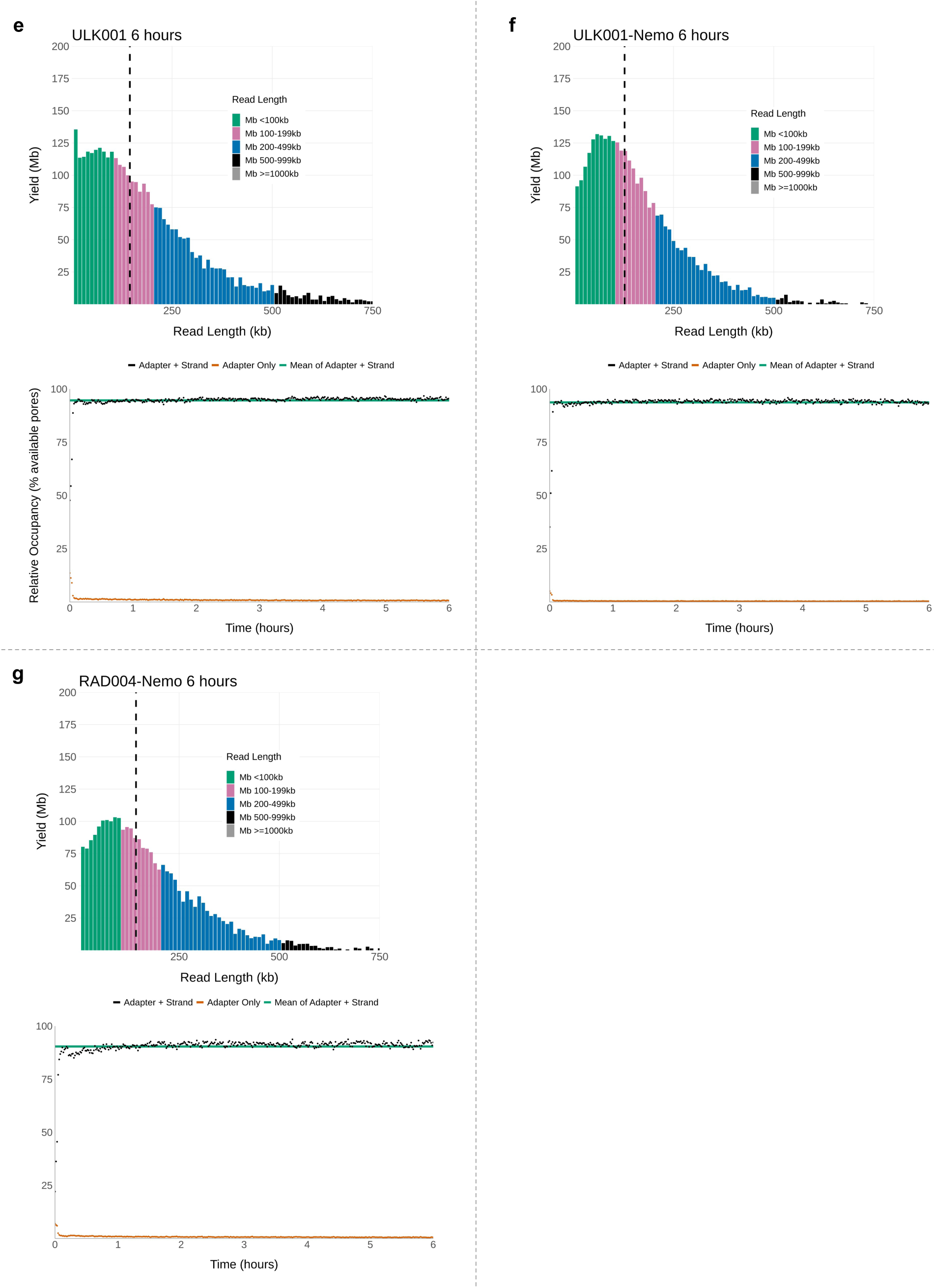
Comparison of sequencing outputs of ultra-long (UL) library preparation protocols. We previously developed two protocols based on the RAD004 kit: Original Quick’s UL v3 (a-b) and KrazyStarFish (KSF) (c-d). New protocols based on Nemo clean-up were developed afterward with comparable outputs to the control ULK001 protocol (e-g). Upper panels = read length distribution with N50 value in black dashed line, lower panels = timelapse of relative pore occupancy. All libraries were prepared from GM12878 cells, loaded on MinION flow cells and sequenced on GridION platform.

**Figure S2.**
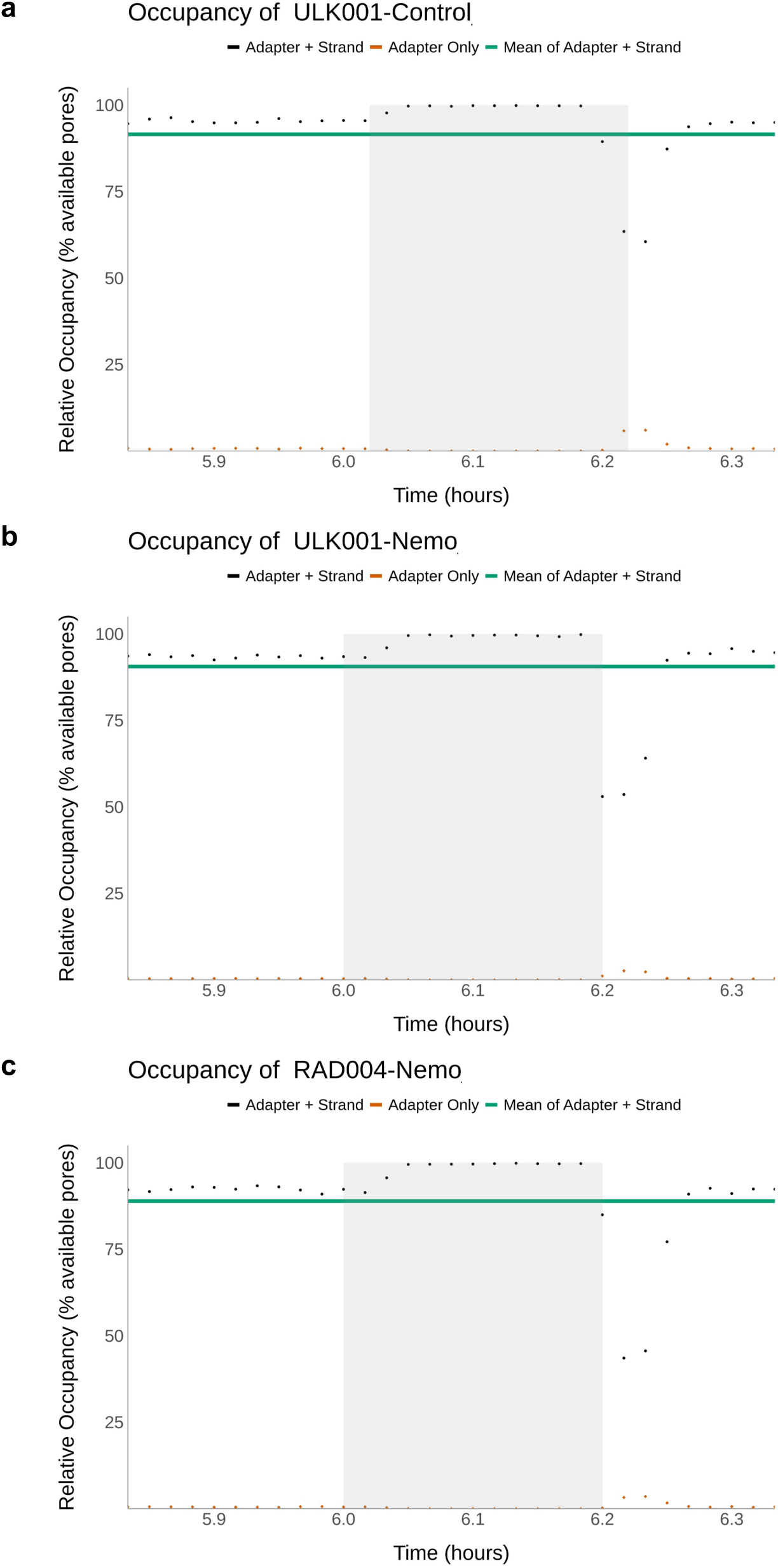
Zoomed-in of occupancy time lapse around mux scan. Due to the “wind-down” time, where non sequencing pores are gradually switched off during the lead into a mux scan, there are time points when relative occupancy reaches 100%

**Figure S3.**
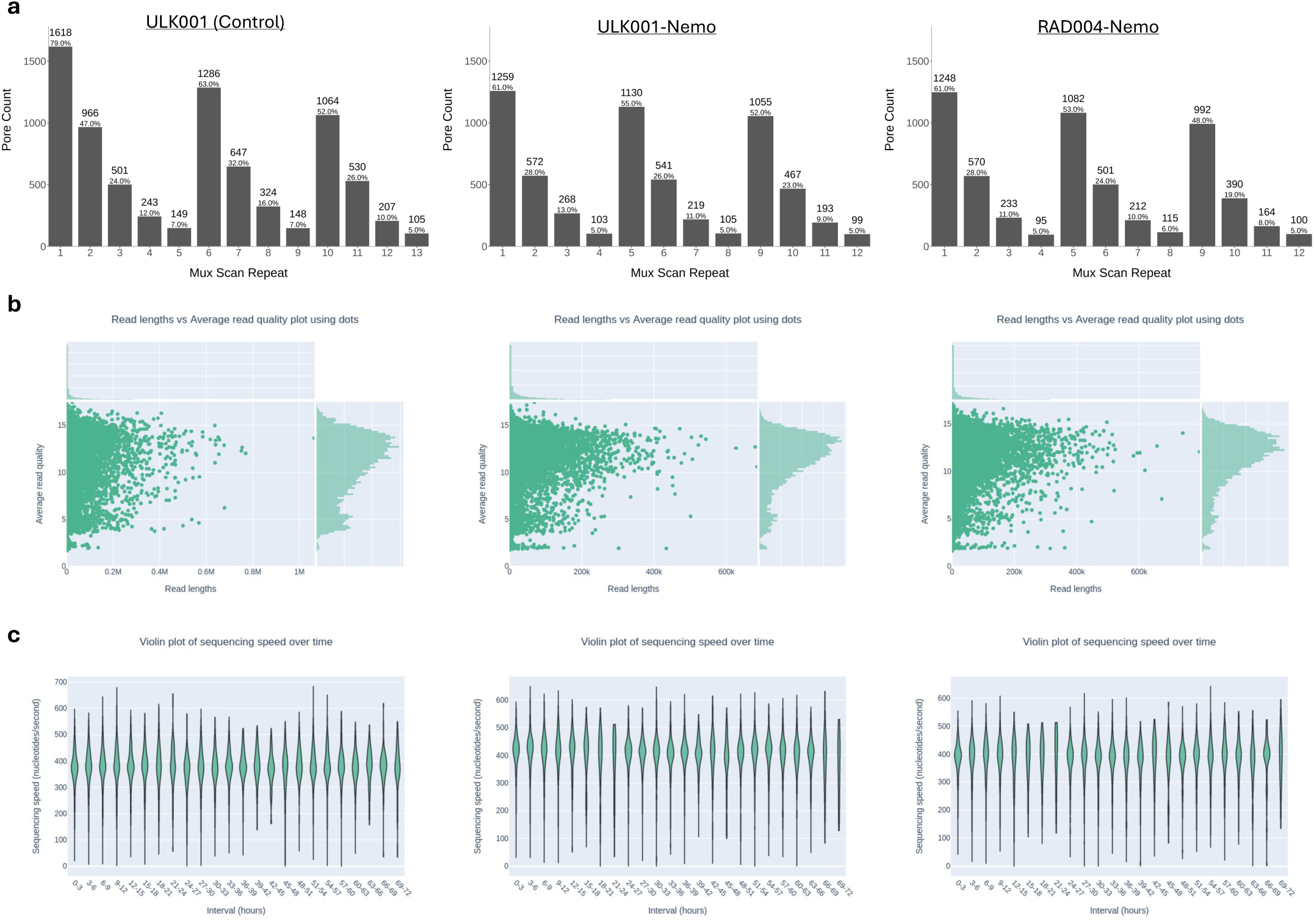
(a) The numbers of available pores after every 6-hour (*i.e.* mux scan) are shown above a percentage of the maximum pore number (i.e. 2048). Length and quality score distributions were similar between the three protocols (b) as well as their sequencing speed (c). Plots (b) and (c) were created by NanoPlot (de Coster 2023). Each loaded library was equivalent to DNA from 3 million GM12878 cells.

**Figure S4.**
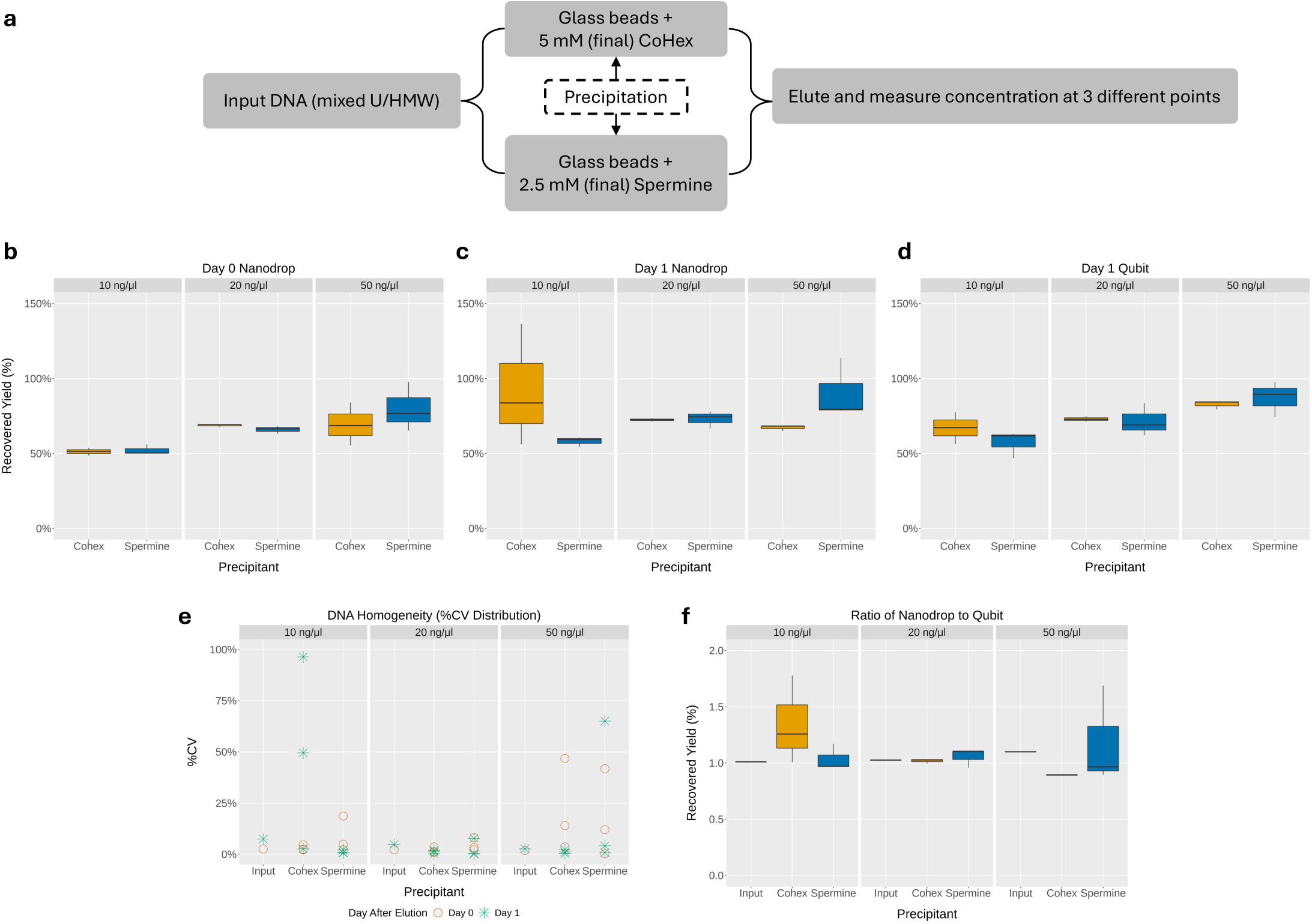
DNA recovery using CoHex and spermine. (a) Workflow of the experiment, (b) Nanodrop measurements of DNA recovery a few hours after elution (Day 0), (c) Nanodrop measurements of DNA recovery after one day (Day 1), (d) Qubit measurements of DNA recovery after one day (Day 1), (e) Homogeneity of samples as measured by %CVs using Nanodrop at Day 0 and Day 1, (f) Ratios of values between Qubit and Nanodrop measurements. DNA was extracted from GM12878 cells and diluted to the respective concentrations (input sample *n*=1, output samples *n*=3).

**Figure S5.**
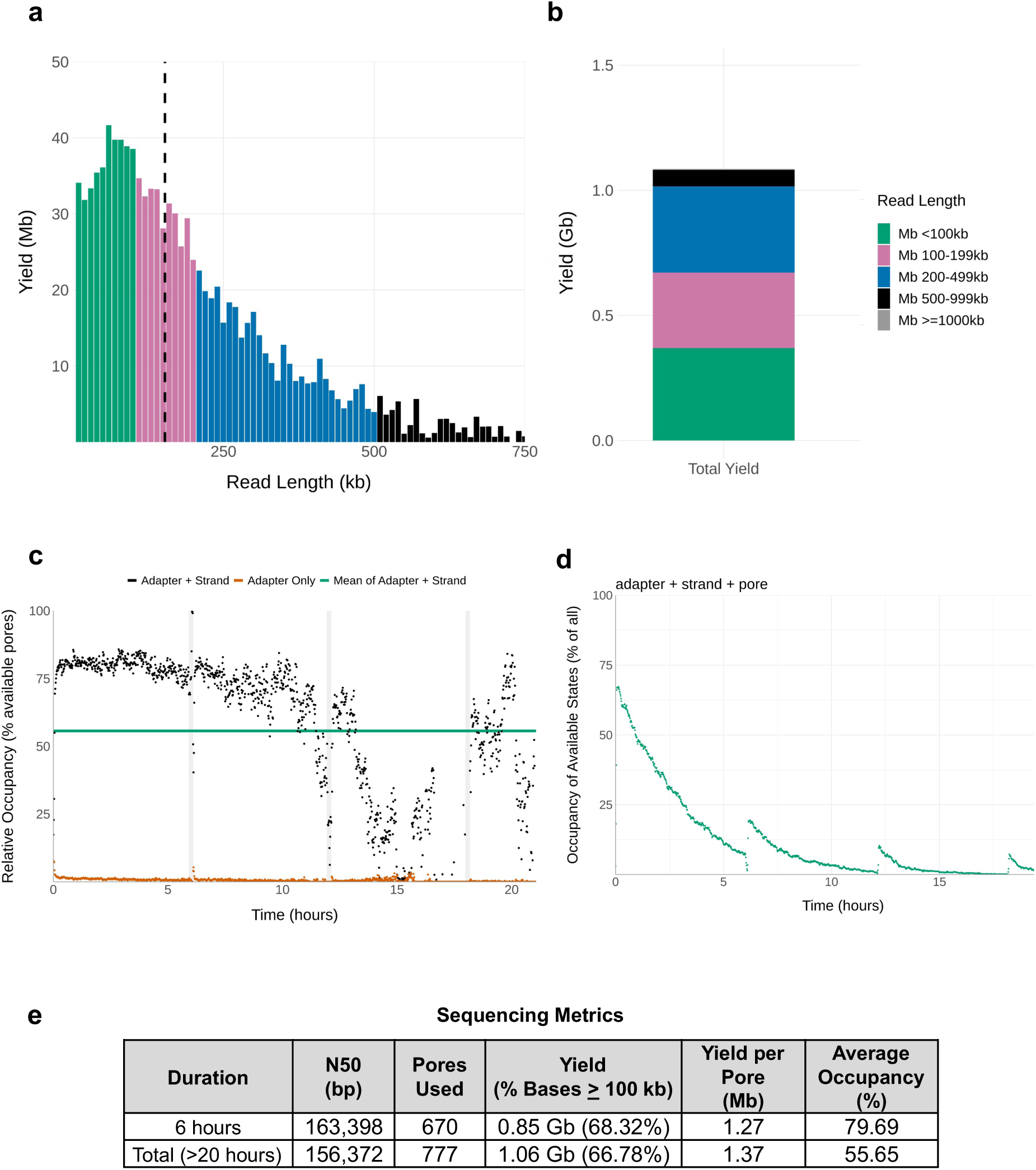
Sequencing output of a concentrated UL library. (a) Read length distribution with N50 marked by the black dashed line, (b) Total yield distribution by read lengths, (c) Time lapse of relative occupancy, (d) Time lapse of available pores showing mux scan at every ∼6 hours, (e) Sequencing metrics are at 6 hours and at the end of data collection (excluding the first 10 minutes). Library was prepared from GM12878 cells, extracted DNA concentration was >60 ng/μl, loaded on a MinION flow cell and sequenced on the GridION platform.

**Figure S6.**
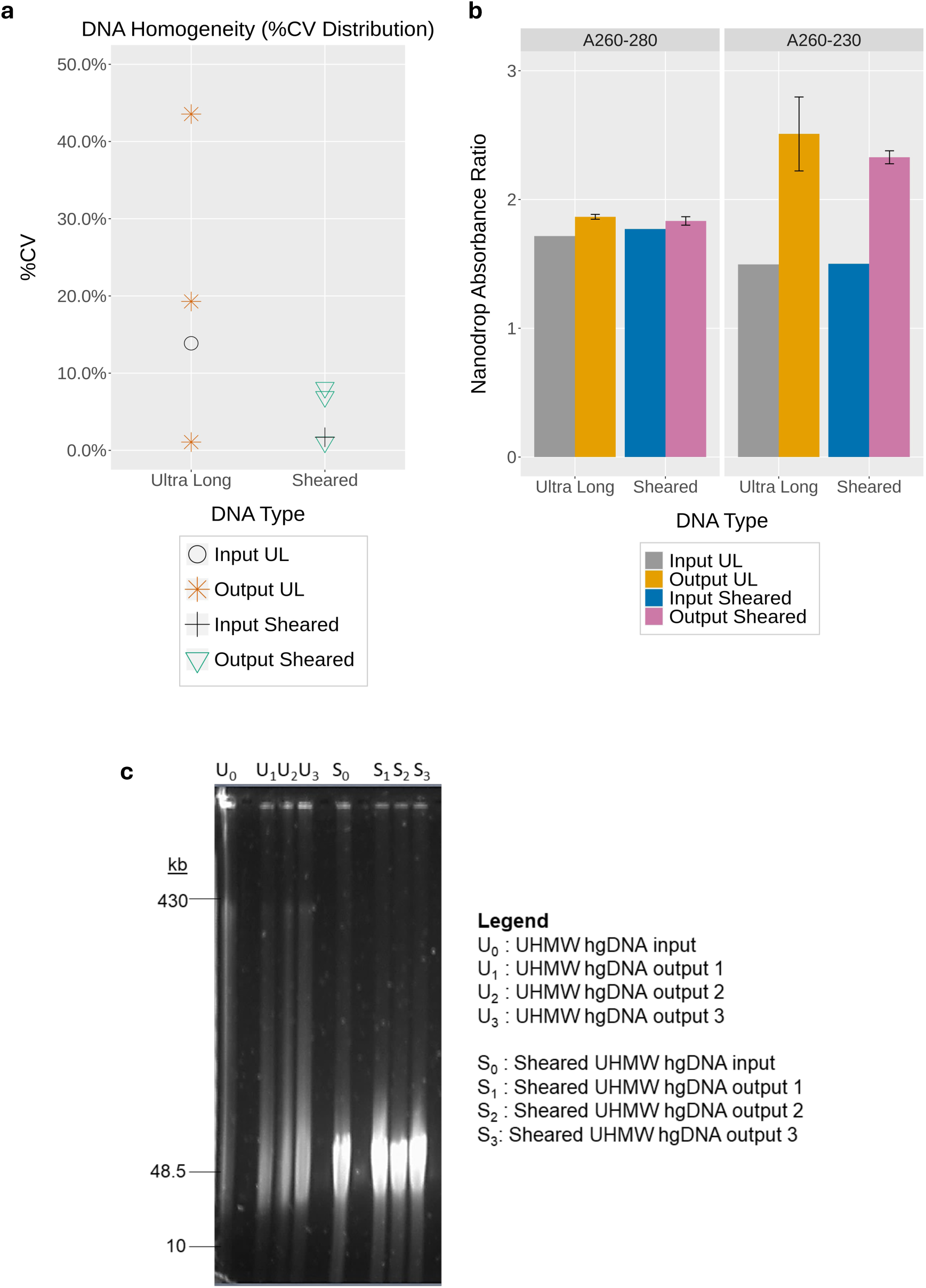
Recovery and quality control of UHMW vs sheared DNA samples by Nemo clean-up. (a) Homogeneity of samples as measured by %CVs using Nanodrop. (b) Absorbance measurements (Nanodrop) of DNA before and after clean-up with Nemo protocol. Nemo clean-up improved the lower quality of input DNA. (c) Pulsed-field gel electrophoresis of samples. Gel concentration was 0.75%, run with Pippin Pulse at the preset mode of ‘5-430 kb’ for 16 hours. UHMW and sheared DNA was extracted from GM12878 cells, diluted to a concentration of ∼15 ng/μl (input sample *n*=1, output samples *n*=3).

**Figure S7.**
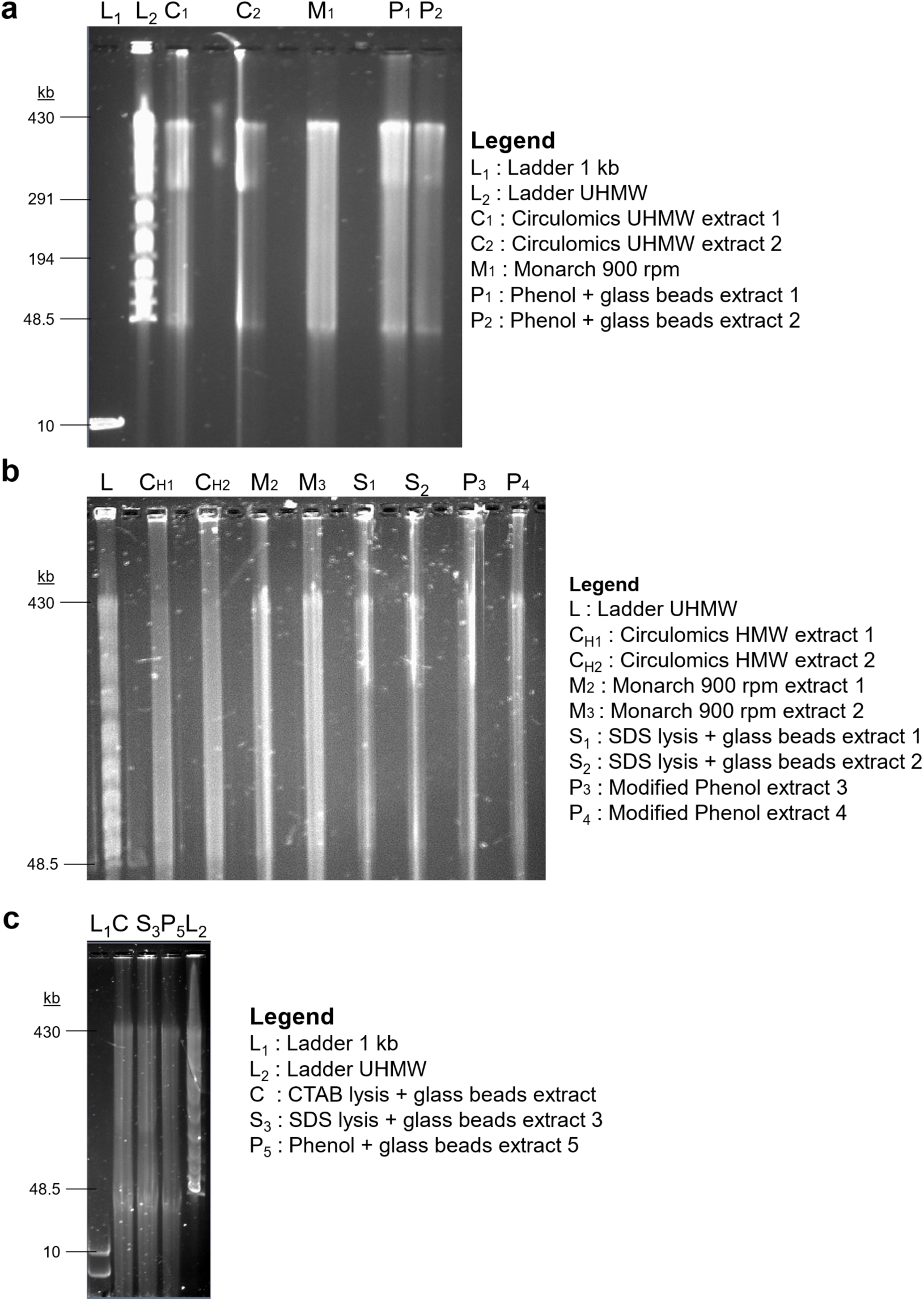
Pulsed-field gel electrophoresis of DNA extracted using different methods, showing their (U)HMW properties. Agarose gel was at 0.75% concentration and was run using Pippin Pulse at the pre-set protocol of “5-430 kb” for 16 hours.

**Figure S8.**
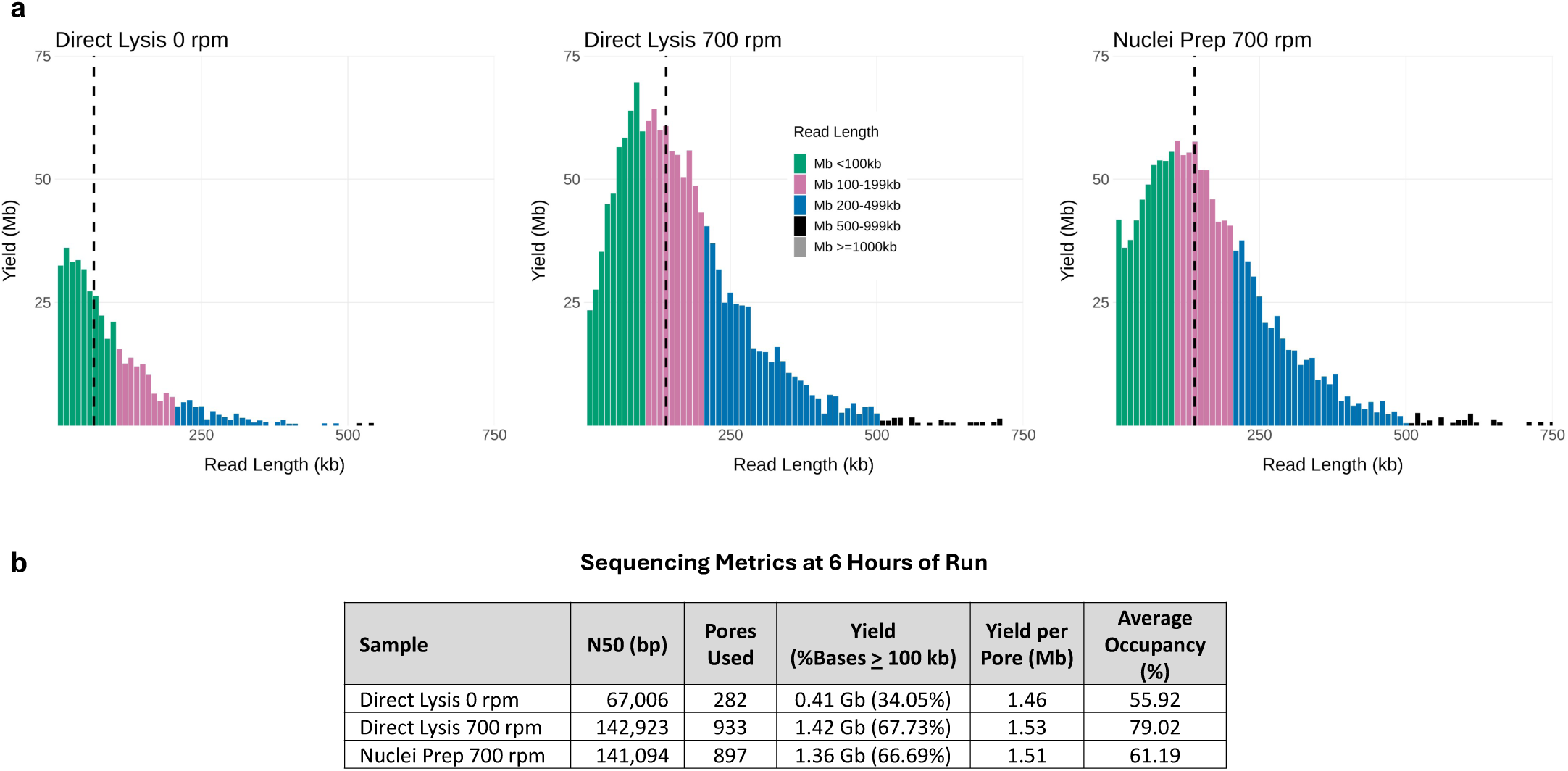
Performance of UL sequencing of libraries extracted with different lysis approaches using Monarch kit, i.e. Direct Lysis vs Nuclei Prep method. Direct lysis approach produced slightly better sequencing metrics than Nuclei Prep. (a) Read length histograms of the runs, (b) Sequencing metrics of the runs at 6 hours (excluding the first 10 minutes). All libraries were prepared from GM12878 cells, loaded on MinION flow cells R9.4 and sequenced on the GridION platform.

**Figure S9.**
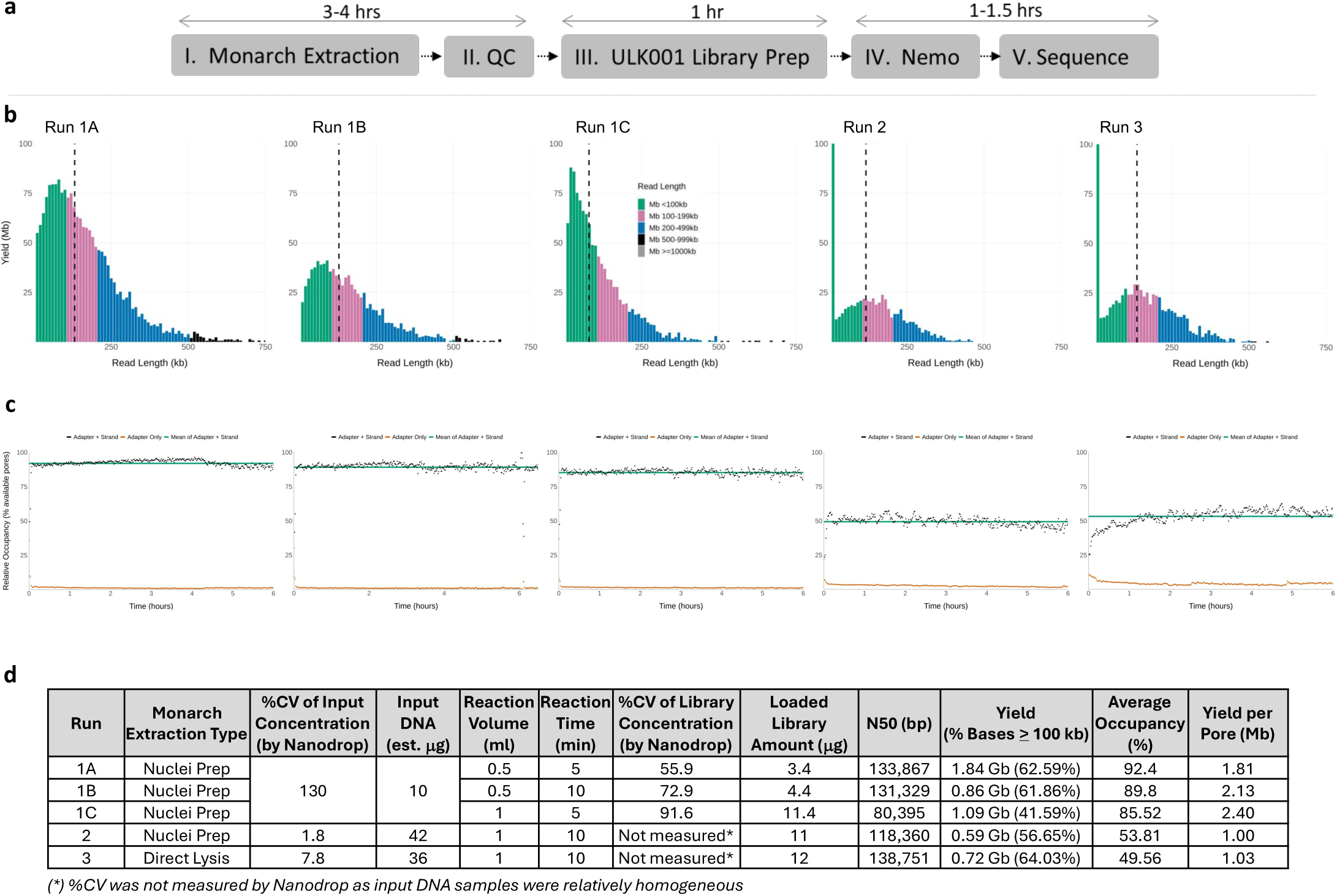
*FindingNemo in one day*: an UL sequencing protocol that extracts UHMW DNA from human cells and sequencing it in a working day (a) The protocol workflow along with the estimated step durations (b) Read length distributions of the libraries extracted using the Monarch kit; dashed black vertical lines denote N50s (c) Occupancy time lapses, (d) DNA and sequencing metrics of the libraries. Samples were extracted from GM12878 cells, each library was loaded on a MinION flow cell and run on the GridION platform. Data are after 6 hours of sequencing (excluding the first 10 minutes).

**Figure S10.**
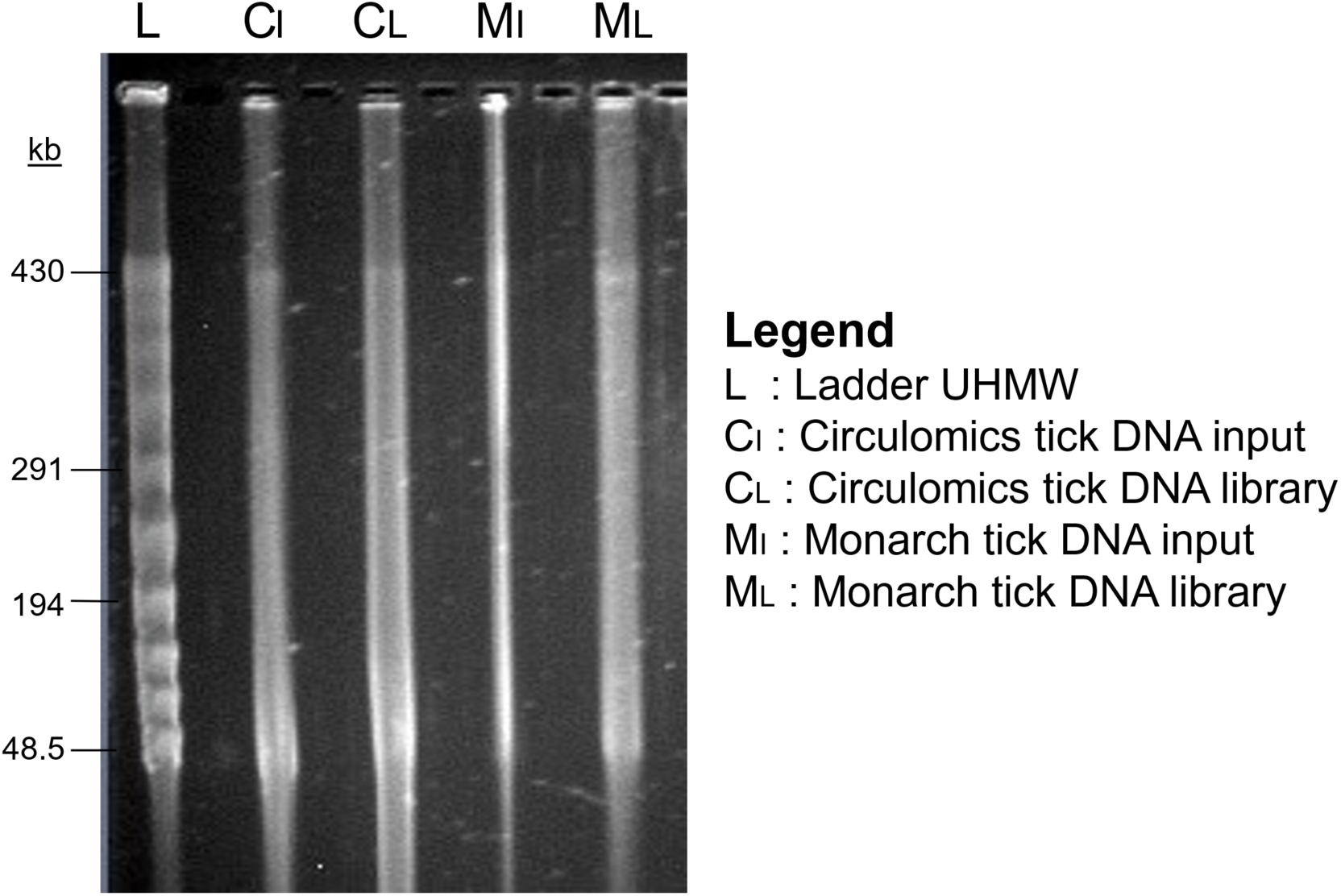
Pulsed-field gel electrophoresis of tick DNA samples prior to and after library preparation based on two different extraction methods: Circulomics and Monarch kit. Agarose gel was at 0.75% concentration and was run using Pippin Pulse at the pre-set protocol “5-430 kb” for 16 hours.

**Table S1.** Sequencing metrics comparison of different library preparation protocols.

| Library | Run Hours (hrs) | N50 (bp) | Pores Used | Yield (% Bases ≥ 100 kb) | Yield per Pore (Mb) | Average Occupancy (%) |
| --- | --- | --- | --- | --- | --- | --- |
| **Previous UL Protocols** |  |  |  |  |  |  |
| A. Quick's UL V3 (1) | 3 | 90,319 | 637 | 0.96 Gb (44.80%) | 1.50 | 78.25 |
| B. Quick's UL V3 (2) | 3 | 85,120 | 1448 | 1.03 Gb (42.60%) | 0.71 | 90.03 |
| C. KrazyStarFish (1) | 6 | 151,704 | 1211 | 1.25 Gb (66.48%) | 1.03 | 66.32 |
| D. KrazyStarFish (2) | 6 | 120,318 | 820 | 1.20 Gb (58.39%) | 1.47 | 66.86 |
| **New UL Protocols** |  |  |  |  |  |  |
| E. ULK001 | 6 | 147,123 | 1025 | 3.27 Gb (64.85%) | 3.19 | 95.01 |
| F. ULK001-Nemo | 6 | 129,678 | 995 | 2.99 Gb (61.71%) | 3.00 | 94.12 |
| G. RAD004-Nemo | 6 | 144,579 | 986 | 2.61 Gb (65.37%) | 2.65 | 90.97 |

**Table S2.**
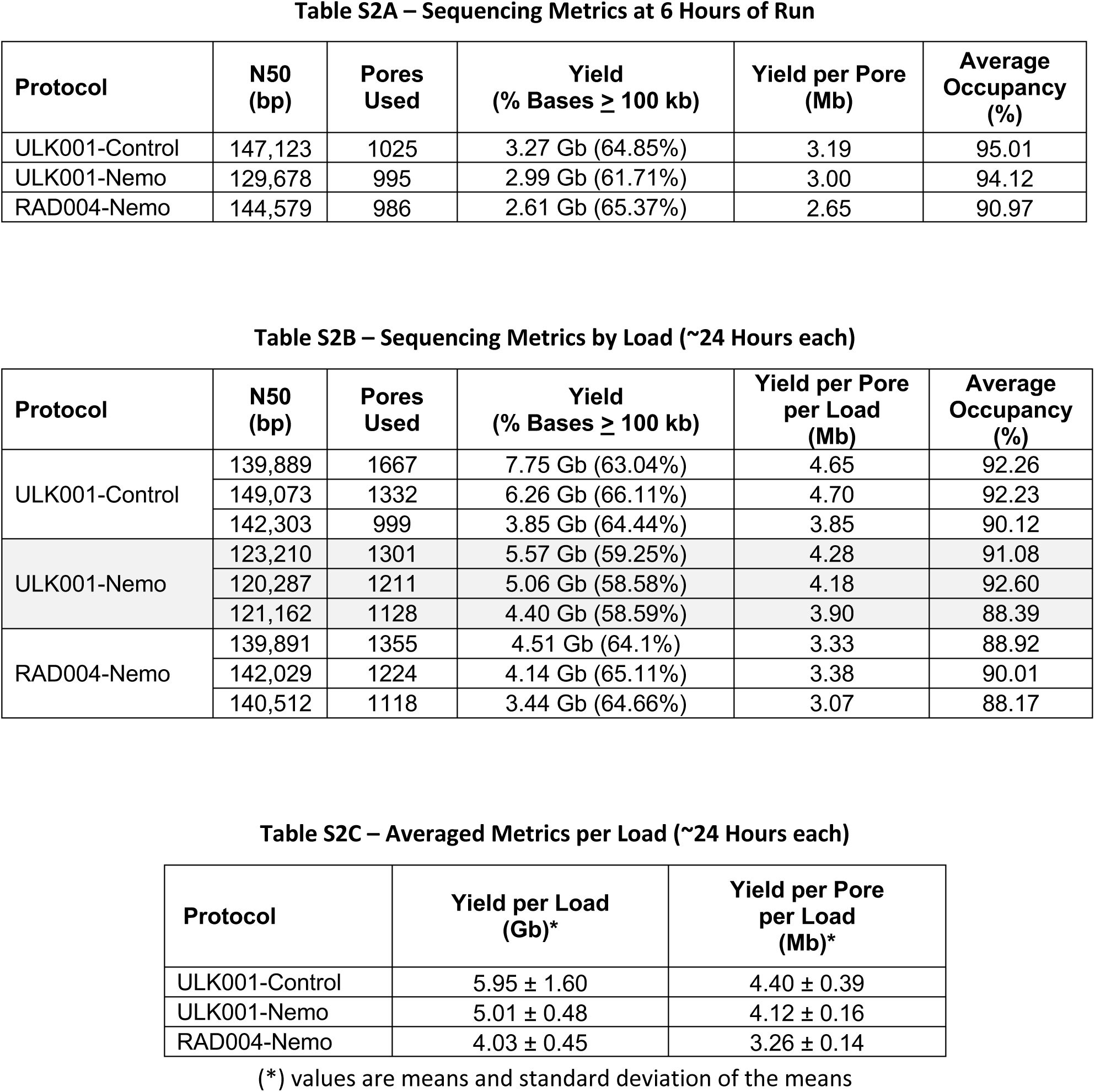
Sequencing Metrics of Rapid-based Kit Libraries.

**Table S3.**
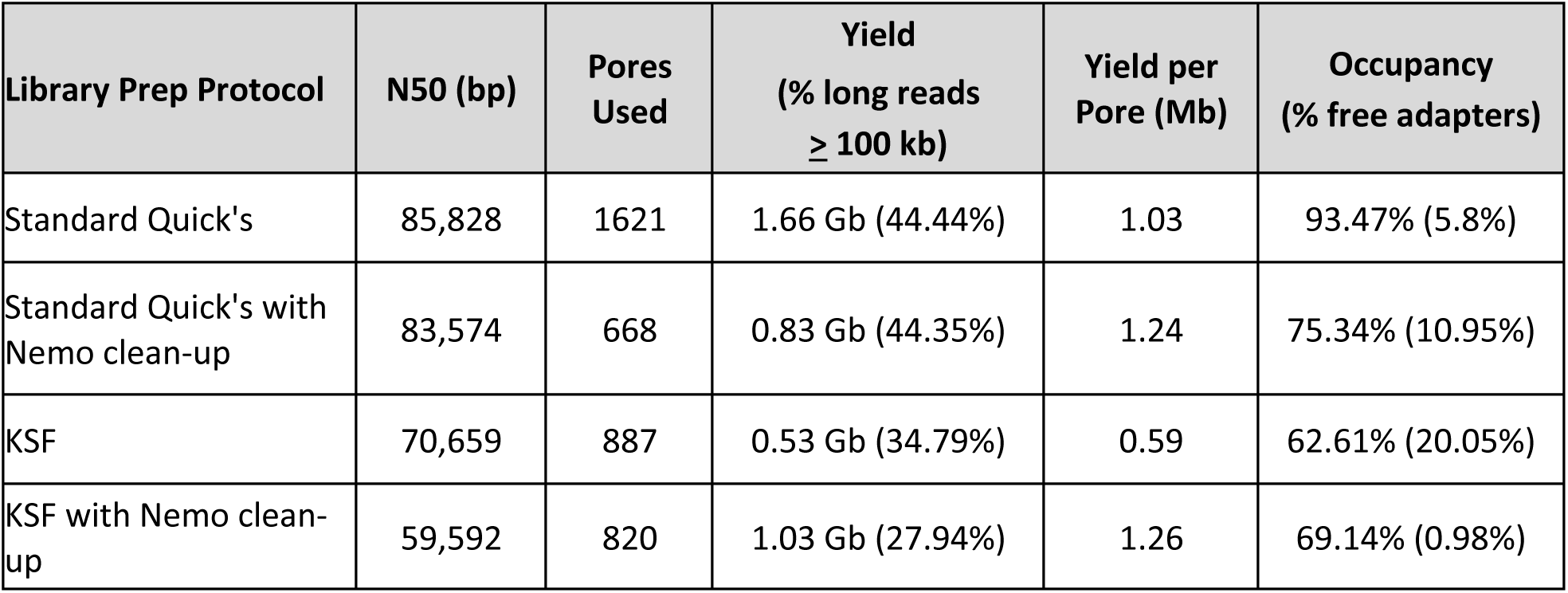
Sequencing metrics of library preparation with or without Nemo clean-up method (4 hours)

| Library Prep Protocol | N50 (bp) | Pores Used | Yield<br>(% long reads<br>≥ 100 kb) | Yield per<br>Pore (Mb) | Occupancy<br>(% free adapters) |
| --- | --- | --- | --- | --- | --- |
| Standard Quick's | 85,828 | 1621 | 1.66 Gb (44.44%) | 1.03 | 93.47% (5.8%) |
| Standard Quick's with<br>Nemo clean-up | 83,574 | 668 | 0.83 Gb (44.35%) | 1.24 | 75.34% (10.95%) |
| KSF | 70,659 | 887 | 0.53 Gb (34.79%) | 0.59 | 62.61% (20.05%) |
| KSF with Nemo clean-up | 59,592 | 820 | 1.03 Gb (27.94%) | 1.26 | 69.14% (0.98%) |

**Table S4.** Library prep conditions and sequencing metrics of samples extracted using the NEB Monarch kit (6 hours)

| Sample | N50 (bp) | Yield<br>(Reads >100 kb) | Average<br>Occupancy<br>(%) | Yield<br>per Pore<br>(Mb) | Library<br>Kit | Preparation<br>Time | Clean-up | Lysis<br>Speed | Preparation<br>Type | Input Cell | Input DNA<br>(µg) | Loaded DNA<br>(µg) |
| --- | --- | --- | --- | --- | --- | --- | --- | --- | --- | --- | --- | --- |
| Monarch-01 | 124,929 | 2.1 Gb (60.18%) | 94.43 | 2.71 | ULK001 | Standard | Nemo | 300 | Nuclei Prep | 1 million | 5 | 2 |
| Monarch-02 | 112,810 | 1.25 Gb (55.99%) | 88.8 | 2.43 | ULK001 | Standard | Nemo | 300 | Nuclei Prep | 1 million | 5 | 1 |
| Monarch-03 | 141,094 | 1.36 Gb (66.69%) | 61.19 | 1.51 | ULK001 | Standard | Nemo | 700 | Nuclei Prep | 3 million | 16 | 9.5 |
| Monarch-04 | 106,493 | 3.35 Gb (52.7%) | 97.36 | 3.89 | ULK001 | Standard | Nanobind | 900 | Nuclei Prep | 3 million | 20 | 18.8 |
| Monarch-05 | 99,327 | 0.56 Gb (49.55%) | 41.59 | 1.06 | RAD004 | Standard | Nemo | 900 | Nuclei Prep | NA | 2 | 1.3 |
| Monarch-06 | 67,006 | 0.41 Gb (34.05%) | 55.92 | 1.46 | ULK001 | Standard | Nemo | 0 | Direct Lysis | 3 million | 20 | 15 |
| Monarch-07 | 142,923 | 1.42 Gb (67.73%) | 79.02 | 1.53 | ULK001 | Standard | Nemo | 700 | Direct Lysis | 3 million | 21 | 5 |
| Monarch-08 | 153,295 | 1.59 Gb (72.55%) | 85.65 | 1.57 | ULK001 | Standard | Spermine<br>(ONT PPT) | 700 | Direct Lysis | 1.5 million | 10 | 1.5 |
| Monarch-09 | 152,653 | 1.61 Gb (72.2%) | 89.29 | 2.03 | ULK001 | Standard | Nemo | 700 | Direct Lysis | 1.5 million | 10 | 3 |
| Monarch-10 | 147,386 | 1.28 Gb (69.51%) | 85.32 | 2.01 | ULK001 | Standard | Spermine +<br>glass beads | 700 | Direct Lysis | 1.5 million | 10 | 5 |
| One Day 1A | 133,867 | 1.84 Gb (62.59%) | 92.4 | 1.81 | ULK001 | One day | Nemo | 600 | Nuclei Prep | 2 million | 10 | 3.5 |
| One Day 1B | 80,395 | 1.09 Gb (41.59%) | 85.52 | 2.40 | ULK001 | One day | Nemo | 600 | Nuclei Prep | 2 million | 10 | 11.5 |
| One Day 1C | 131,329 | 0.86 Gb (61.86%) | 89.8 | 2.13 | ULK001 | One day | Nemo | 600 | Nuclei Prep | 2 million | 10 | 4.5 |
| One Day 2 | 118,360 | 0.59 Gb (56.65%) | 53.81 | 1.00 | ULK001 | One day | Nemo | 800 | Nuclei Prep | 3 million | 42 | 11 |
| One Day 3 | 138,751 | 0.72 Gb (64.03%) | 49.56 | 1.03 | ULK001 | One day | Nemo | 800 | Direct Lysis | 3 million | 36 | 12 |

**Table S5.** Comparison of runs on MinION vs PromethION platform.

| Sample | Platform | N50 (bp) | Pore No. | Yield<br>(Reads >100 kb) | Yield per<br>Pore (Mb) | Average<br>Occupancy<br>(%) |
| --- | --- | --- | --- | --- | --- | --- |
| Tick-Circulomics | MinION | 109,486 | 527 | 1.55 Gb (52.98%) | 2.94 | 80.11 |
| Tick-Circulomics | PromethION | 104,993 | 3492 | 11.11 Gb (51.64%) | 3.18 | 87.50 |
| Tick-Monarch | MinION | 123,683 | 617 | 1.85 Gb (60.25%) | 2.99 | 84.71 |
| Tick-Monarch | PromethION | 126,008 | 3902 | 15.01 Gb (61.46%) | 3.85 | 88.20 |

## References

Allers T, Lichten M. 2000. A method for preparing genomic DNA that restrains branch migration of Holliday junctions. Nucleic Acids Res 28: 6.

Arscott PG, Ma C, Wenner JR, Bloomfield VA. 1995. DNA condensation by cobalt hexammine(III) in alcohol–water mixtures: Dielectric constant and other solvent effects. Biopolymers 36: 345–364. https://onlinelibrary.wiley.com/doi/10.1002/bip.360360309 (Accessed February 25, 2022).

Brekke, T.D., Papadopulos, A.S.T., Julià, E., Fornas, O., Fu, B., Yang, F., de la Fuente, R., Page, J., Baril, T., Hayward, A., Mulley, J.F., 2023. A New Chromosome-Assigned Mongolian Gerbil Genome Allows Characterization of Complete Centromeres and a Fully Heterochromatic Chromosome. Mol Biol Evol 40. 10.1093/molbev/msad115.

Cahyani I, Tyson J, Holmes N, Quick J, Loman N, Loose M. 2021a. FindingNemo: A Toolkit of CoHex- and Glass Bead-based Protocols for Ultra-Long Sequencing on ONT Platforms. protocols.io. https://www.protocols.io/view/findingnemo-a-toolkit-of-cohex-and-glass-bead-base-dm6gpwkn1lzp/v1 (Accessed March 11, 2024).

Cahyani I, Tyson J, Holmes N, Quick J, Loman N, Loose M. 2021b. FindingNemo in OneDay: Ultra-Long ONT Library Preparation from Cell to Flow Cell in One Day. protocols.io.10.17504/protocols.io.bxhxpj7n

Cahyani I, Tyson J, Holmes N, Quick J, Loman N, Loose M. 2024. FindingNemo (v.kit14): A Toolkit for DNA Extraction, Library Preparation and Purification for Ultra Long Nanopore Sequencing. protocols.io. 10.17504/protocols.io.5jyl8p38rg2w/v1

Cahyani, I., 2024. Ultra-Long Sequencing of Yeast Cells (S. cerevisiae) on ONT Sequencers – A Modified FindingNemo Protocol. protocols.io. URL https://www.protocols.io/view/ultra-long-sequencing-of-yeast-cells-s-cerevisiae-rm7vzjbk5lx1/v1 (accessed 5.16.24).

de Coster W, Rademakers R. 2023. NanoPack2: population-scale evaluation of long-read sequencing data ed. C. Alkan. Bioinformatics 39. https://academic.oup.com/bioinformatics/article/doi/10.1093/bioinformatics/btad311/7160911 (Accessed March 11, 2024).

Deng H, Bloomfield VA. 1999. Structural effects of cobalt-amine compounds on DNA condensation. Biophys J 77: 1556. /pmc/articles/PMC1300443/?report=abstract (Accessed February 25, 2022).

Giordano F, Aigrain L, Quail MA, Coupland P, Bonfield JK, Davies RM, Tischler G, Jackson DK, Keane TM, Li J, et al. 2017. De novo yeast genome assemblies from MinION, PacBio and MiSeq platforms. Sci Rep 7: 1–10.

‘Giron’ Koetsier PA, Cantor EJ 2021. A simple approach for effective shearing and reliable concentration measurement of ultra-high-molecular-weight DNA. Biotechniques 71: 439–444. https://www.future-science.com/doi/10.2144/btn-2021-0051 (Accessed August 4, 2021).

Jain M, Koren S, Miga KH, Quick J, Rand AC, Sasani TA, Tyson JR, Beggs AD, Dilthey AT, Fiddes IT, et al. 2018a. Nanopore sequencing and assembly of a human genome with ultra-long reads. Nat Biotechnol 36: 338–345. https://www.nature.com/articles/nbt.4060.

Jain, M., Olsen, H., Turner, D. et al. 2018b. Linear assembly of a human centromere on the Y chromosome. Nat Biotechnol 36, 321–323. 10.1038/nbt.4109

Kankia BI, Buckin V, Bloomfield VA. 2001. Hexamminecobalt(III)-induced condensation of calf thymus DNA: circular dichroism and hydration measurements. Nucleic Acids Res 29: 2795. /pmc/articles/PMC55774/ (Accessed July 19, 2022).

Koetsier G, Cantor E. 2019. A Practical Guide to Analyzing Nucleic Acid Concentration and Purity with Microvolume Spectrophotometers. New Engl Biolabs 1: 1–8. https://www.neb.com/-/media/catalog/application-notes/mvs_analysis_of_na_concentration_and_purity.pdf?rev=be7c8e19f4d34e558527496ea51623dc.

Lin Y-C, Boone M, Meuris L, Lemmens I, Roy N Van, Soete A, Reumers J, Moisse M, Plaisance S, Drmanac R, et al. 2014. Genome dynamics of the human embryonic kidney 293 lineage in response to cell biology manipulations. Complet. Genomics Inc 120: 12. www.nature.com/naturecommunications (Accessed March 11, 2024).

Logsdon GA, Vollger MR, Hsieh PH, Mao Y, Liskovykh MA, Koren S, Nurk S, Mercuri L, Dishuck PC, Rhie A, et al. 2021. The structure, function and evolution of a complete human chromosome 8. Nature. 10.1038/s41586-021-03420-7.

LongRead Club, n.d. Long Read Club. protocols.io. https://www.protocols.io/workspaces/long-read-club/about (accessed 5.15.24).

Miga KH. 2020. Centromere studies in the era of ‘telomere-to-telomere’ genomics. Exp Cell Res 394: 112127. 10.1016/j.yexcr.2020.112127.

NEB Direct Lysis. Protocol Guidance for Extraction of Ultra-High Molecular Weight (UHMW) Genomic DNA for Ultra-Long (UL) Read NGS Sequencing applications in Oxford Nanopore Technologies® workflows. New Engl Biolabs. https://www.neb.com/en-gb/protocols/2022/02/17/protocol-guidance-for-extraction-of-high-molecular-weight-uhmw-genomic-dna-for-ultra-long-ul-read-ngs-sequencing-applications-in-oxford-nanopore-technologies-workflows (Accessed March 12, 2024).

NEB T3050. INSTRUCTION MANUAL Monarch ® HMW DNA Extraction Kit for Cells & Blood. New Engl Biolabs. www.neb.com/T3050 (Accessed March 12, 2024).

ONT ULK001. Ultra-Long Sequencing Kit (SQK-ULK001) Protocol. Nanopore Community. https://community.nanoporetech.com/docs/prepare/library_prep_protocols/μltra-long-sequencing-kit-ULK001/v/μlk_9124_v110_revn_24mar2021 (Accessed March 12, 2024).

ONT ULK114. Ultra-Long DNA Sequencing Kit V14 (SQK-ULK114) Protocol. Nanopore Community. https://community.nanoporetech.com/docs/prepare/library_prep_protocols/μltra-long-dna-sequencing-kit-sqk-ulk114/v/μlk_9177_v114_revl_27nov2022 (accessed 5.15.24).

Ouameur AA, Tajmir-Riahi H-A. 2004. Structural Analysis of DNA Interactions with Biogenic Polyamines and Cobalt(III)hexamine Studied by Fourier Transform Infrared and Capillary Electrophoresis. J Biol Chem 279: 42041–42054. https://linkinghub.elsevier.com/retrieve/pii/S002192582072675X (Accessed February 25, 2022).

PacBio, 2023. Nanobind ® CBB kit For extraction of HMW genomic DNA from cultured cells, cultured bacteria, and blood. [WWW Document]. URL https://www.pacb.com/wp-content/uploads/Guide-overview-Nanobind-CBB-kit.pdf (accessed 5.15.24).

Pelta J, Livolant F, Sikorav J-L. 1996. DNA Aggregation Induced by Polyamines and Cobalthexamine. J Biol Chem 271: 5656–5662. https://linkinghub.elsevier.com/retrieve/pii/S0021925818979068 (Accessed March 11, 2024).

Quick J. 2018. Ultra-long read sequencing protocol for RAD004. protocols.io. 10.17504/protocols.io.mrxc57n.

Robertson RM, Smith DE. 2007. Strong effects of molecular topology on diffusion of entangled DNA molecules. Proc Natl Acad Sci U S A 104: 4824–4827. www.pnas.org/cgidoi10.1073pnas.0700137104 (Accessed March 12, 2024).

Rodrigue S, Materna AC, Timberlake SC, Blackburn MC, Malmstrom RR, Alm EJ, Chisholm SW. 2010. Unlocking Short Read Sequencing for Metagenomics. PLoS One 5: e11840. https://journals.plos.org/plosone/article?id=10.1371/journal.pone.0011840 (Accessed February 7, 2022).

Salazar AN, Gorter de Vries AR, van den Broek M, Wijsman M, de la Torre Cortés P, Brickwedde A, Brouwers N, Daran J-MG, Abeel T. 2017. Nanopore sequencing enables near-complete de novo assembly of Saccharomyces cerevisiae reference strain CEN.PK113-7D. FEMS Yeast Res 17. https://academic.oup.com/femsyr/article/17/7/fox074/4157789 (Accessed July 22, 2021).

sigmaaldrich.com. Spermine. https://www.sigmaaldrich.com/deepweb/assets/sigmaaldrich/product/documents/122/740/s3256pis.pdf (Accessed March 11, 2024).

Simbolo M, Gottardi M, Corbo V, Fassan M, Mafficini A. 2013. DNA Qualification Workflow for Next Generation Sequencing of Histopathological Samples. PLoS One 8: 62692. www.plosone.org (Accessed March 12, 2024).

Stortchevoi A, Kamelamela N, Levine SS. 2020. SPRI beads-based size selection in the range of 2-10kb. J Biomol Tech 31: 7–10. /pmc/articles/PMC6944320/ (Accessed February 7, 2022).

Synthego HEK293. HEK293 Cells: Background, Applications, Protocols, and More. synthego.com. https://www.synthego.com/hek293 (Accessed March 12, 2024).

